# 5’ UTR variants in the quantitative trait gene *Hnrnph1* support reduced 5’ UTR usage and hnRNP H protein as a molecular mechanism underlying reduced methamphetamine sensitivity

**DOI:** 10.1101/2020.01.11.902908

**Authors:** Qiu T. Ruan, Neema Yazdani, Eric R. Reed, Jacob A. Beierle, Lucy P. Peterson, Kimberly P. Luttik, Karen K. Szumlinski, William E. Johnson, Peter E. A. Ash, Benjamin Wolozin, Camron D. Bryant

## Abstract

We previously identified a 210 kb region on chromosome 11 (50.37-50.58 Mb, mm10) containing two protein-coding genes (*Hnrnph1, Rufy1*) that was necessary for reduced methamphetamine-induced locomotor activity in C57BL/6J congenic mice harboring DBA/2J polymorphisms. Gene editing of a small deletion in the first coding exon supported *Hnrnph1* as a quantitative trait gene. We have since shown that *Hnrnph1* mutants also exhibit reduced methamphetamine-induced reward, reinforcement, and dopamine release. However, the quantitative trait variants (**QTVs**) that modulate *Hnrnph1* function at the molecular level are not known. Nine single nucleotide polymorphisms and seven indels distinguish C57BL/6J from DBA/2J within *Hnrnph1*, including four variants within the 5’ untranslated region **(UTR)**. Here, we show that a 114 kb introgressed region containing *Hnrnph1* and *Rufy1* was sufficient to cause a decrease in MA-induced locomotor activity. Gene-level transcriptome analysis of striatal tissue from 114 kb congenics versus *Hnrnph1* mutants identified a nearly perfect correlation of fold-change in expression for those differentially expressed genes that were common to both mouse lines, indicating functionally similar effects on the transcriptome and behavior. Exon-level analysis (including noncoding exons) revealed decreased 5’ UTR usage of *Hnrnph1* and immunoblot analysis identified a corresponding decrease in hnRNP H protein in 114 kb congenic mice. Molecular cloning of the *Hnrnph1* 5’ UTR containing all four variants (but none of them individually) upstream of a reporter induced a decrease in reporter signal in both HEK293 and N2a cells, thus identifying a set of QTVs underlying molecular regulation of *Hnrnph1*.

## INTRODUCTION

Psychostimulant use disorders (**PUDs**), including methamphetamine (**MA**) and cocaine dependence, are a serious public health concern in the United States. While the opioid epidemic crisis continues to garner warranted attention, there has been much less focus on the recent steep surge in PUDs as evidenced by the steep increase in PUD-related deaths, especially MA-related deaths (1). This public health concern is particularly problematic, given that there are no FDA-approved treatments for PUDs. Both genetic and environmental factors contribute to PUDs (2), yet genome-wide association studies to date have identified very few genetic factors (3). One notable example was the identification of a genome-wide association between *FAM53B* and cocaine dependence in humans (4). This finding was particularly interesting in the context of a mouse forward genetic study of psychostimulant addiction traits that identified a *trans*-expression quantitative trait locus (**QTL**) originating from a locus containing *Cyfip2* and *Hnrnph1* that influenced *Fam53b* expression and was associated with cocaine intravenous self-administration (5). This is one of the first examples demonstrating a direct correspondence between genome-wide genetic association, functional changes in gene expression, and trait-relevant behaviors (cocaine self-administration) between rodents and humans. Thus, this example highlights the power and translational relevance of systems genetics in mice in the study of substance use disorder (**SUD)**-relevant traits.

There are several advantages to using rodent forward genetics in studying the genetic basis of molecular and behavioral traits relevant to SUDs, including the ability to control allelic frequency, genetic background, the environment, the amount and length of drug exposure, the sample size, the collection of the appropriate tissue at the appropriate time points, and the ability to both map and validate causal functional variants *in vivo* within the same animal species. While there are clear limitations to modeling the genetic basis of psychiatric disease traits in rodents, it is important to point out studies in which there have been promising successes. In addition to the translational link of *Fam53b* with cocaine addiction traits in mice and humans (4, 5), there are multiple other recent examples of forward genetic discoveries in rodents that have yielded a high likelihood for translation to humans. A missense SNP in *Taar1* (trace amine-associated receptor 1) was mapped in mice for the aversive properties of MA self-administration, body temperature (6) and toxicity (7–9) and genetic variants in *TAAR1* in humans were associated with drug craving in methamphetamine dependence (10, 11). *CNIH3* (Cornichon Family AMPA Receptor Auxiliary Protein 3) was identified as a GWAS hit for opioid dependence in humans that was corroborated with a forward genetic study of morphine physical dependence in mice (12). Whole genome sequencing plus QTL analysis and functional validation identified *Grm2* (mGlu2 receptor) in escalation of alcohol consumption in rats (13). A mouse strain survey correlating gene expression with behavior found a link between glyoxylase 1 (*Glo1*) and anxiety-like behavior that was confirmed via viral knockdown and overexpression (14). A *cis*-eQTL and copy number variant containing *Glo1* was subsequently discovered to segregate in outbred mice and is fixed in several classical inbred strains, thus revealing a genetic and molecular mechanism linking *Glo1* expression with anxiety-like behavior (15). *Glo1* metabolizes methylglyoxal which, in light of mouse preclinical studies, was found to act as an agonist for the GABA-A receptor and exert anxiolytic-like effects (16, 17). Increasing methylglyoxal levels preclinically reduced alcohol drinking (18, 19), induced a rapid antidepressant-like response (20), and reduced seizures (21). To summarize, mouse forward genetic studies have led to the identification and/or corroboration of several promising therapeutic targets for SUDs and other psychiatric disorders.

A major focus of our lab is to use discovery-based forward genetics to identify novel genetic factors underlying heritable differences in sensitivity to the locomotor stimulant response to drugs of abuse, including opioids as well as psychostimulants such as MA. The neurocircuitry and neurochemical mechanisms underlying drug-induced locomotor activity are in part shared with those that mediate the addictive properties of drugs of abuse (22–24); thus, a reasonable hypothesis is that a subset of shared, polymorphic genetic factors influence both sets of complex behavioral traits. Examples from our lab that support this hypothesis include casein kinase 1-epsilon (***Csnk1e***) which influences opioid- and MA-induced locomotor activity (25, 26) and opioid conditioned place preference (27) as well as heterogeneous nuclear ribonucleoprotein H1(***Hnrnph1***) which we first mapped and validated for MA-induced locomotor activity (28) and subsequently found influences MA-induced dopamine release in the nucleus accumbens, MA-induced reward, and MA-induced reinforcement (29).

With regard to *Hnrnph1*, QTL mapping in an F2 cross first identified a locus containing *Hnrnph1* on chromosome 11 whereby the DBA/2J (**D2J**) allele was associated with reduced MA-induced locomotor activity (30). An interval-specific congenic approach was employed in C57BL/6J (**B6J**) mice carrying various introgressed regions on chromosome 11 from the DBA/2J strain (31) to fine map the QTL. We detected a fortuitous recombination event that revealed a 210 kb region on chromosome 11 (50.37 - 50.58 Mb) that was necessary for reduced sensitivity in the locomotor stimulant response to MA (28). Replacement of this polymorphic region with the background C57BL/6J allele completely eliminated the MA-induced behavioral phenotype, thus demonstrating that this region was necessary for reduced MA-induced locomotor activity (28). The 210 kb region contains two protein-coding genes – *Hnrnph1* and *Rufy1*. Introduction of a heterozygous deletion within the first coding exon of each gene provided strong support for *Hnrnph1* (and not *Rufy1*) as a quantitative trait gene (**QTG**) underlying reduced MA sensitivity (28) and we subsequently expanded the phenotypic repertoire of the *Hnrnph1* mutants to include a reduction MA reinforcement, MA reward and MA-induced dopamine release (32).

Although the combined published evidence supports *Hnrnph1* as a QTG for MA sensitivity, to date, we have only demonstrated that inheritance of the 210 kb region polymorphic region containing *Hnrnph1* and *Rufy1* is *necessary* for the reduction in MA-induced behavior. In order to demonstrate that this region is also *sufficient*, in the present study, we backcrossed and screened for congenic mice capturing only *Hnrnph1* and *Rufy1* and we identified a founder containing a 114 kb introgressed region. Following the observation of reduced MA-induced locomotor activity in 114 kb congenic mice, we assessed the striatal transcriptome at both the gene- and exon-level and compared these results to the transcriptome of *Hnrnph1* mutants (**H1**^**+/-**^) in order to provide further support for *Hnrnph1* as a QTG and to identify the quantitative trait variant(s) (**QTV**)s. Upon discovering decreased usage of the 5’ UTR in *Hnrnph1* in 114 kb congenic mice as a potential molecular mechanism underlying the QTL and QTG, we validated decreased 5’ UTR expression via real-time quantitative PCR (**qPCR**) by targeting the adjacent exon junction and the specific 5’ UTR noncoding exon. We then identified decreased protein expression of hnRNP H in 114 kb congenic mice as a potential downstream functional consequence of reduced 5’ UTR usage. Finally, to validate candidate QTV(s), we cloned the 5’ untranslated region (**5’ UTR**) of *Hnrnph1* containing either the individual *Hnrnph1* 5’ UTR variants, or the combined set of all four 5’ UTR variants fused to a luciferase reporter gene and tested the effect on reporter expression in two different cell lines. The results identify a set of 5’ UTR variants within *Hnrnph1* that likely represent the QTVs underlying molecular regulation of *Hnrnph1* and behavior.

## MATERIALS AND METHODS

### Generation of B6J.D2J 114 kb congenic mice

All procedures involving mice were conducted in accordance with the Guidelines for Ethical Conduct in the Care and Use of Animals and were approved by the Institutional Animal Care and Use Committee at Boston University (#AN-15326). The founder 114 kb congenic mouse was identified by monitoring the distal single nucleotide polymorphism (**SNP**) sites of offspring generated via backcrossing heterozygous congenic mice from Line 4a carrying one copy of a ∼11 Mb introgressed interval from the D2J strain to the background B6J strain (28). We monitored for retention of the most proximal D2J SNP that was just upstream of *Hnrnph1* (rs29383600; 50,373,006 bp; mm10) and this SNP defined the proximal end of the 114 kb interval. We previously genotyped several purported SNP markers proximal to rs29833600 that were monomorphic until 49,873,463 bp (mm10), which was catalogued by Sanger as a B6J/D2J polymorphic marker and that we genotyped as homozygous for B6J (28). Because we were unable to identify any polymorphisms between 49,873,463 bp (mm10) and rs29383600, we defined the proximal interval of this smaller congenic with the marker rs29383600 (50,373,006 bp). The remainder of the genome is isogenic for B6J as determined by several other markers on chromosome 11 and by a medium-density genotyping array containing 882 informative SNP markers from our previous study (28).

We also simultaneously monitored for the detection of a recombination event (i.e., the observation of a homozygous B6J genotype) at a second marker located just distal to *Rufy1* (rs254771403; 50,486,998 bp; mm10). Following detection of a recombination event at rs254771403, we then genotyped at an additional upstream proximal marker at rs29459915 (50,484,260 bp; mm10) that was also downstream of *Rufy1* and found retention of the D2J allele at this locus. Thus, the new remarkably smaller congenic interval was conservatively defined by a region spanning proximal rs29383600 (50,373,006 bp) and distal rs254771403 (50,486,498 bp), yielding a 114 kb interval. Thus, we named this new congenic, **“114 kb”**.

PCR primers were designed to amplify a ∼200 bp amplicon that contained and flanked each SNP. The PCR products were run on a 1.5% agarose gel and visualized for band specificity. Single bands were excised according to their predicted fragment size and gel-purified (Promega Wizard SV Gel and PCR Clean-Up System, Cat A9281), and prepared for Sanger sequencing (Genewiz, Cambridge, MA, USA). Mice homozygous for the 114 kb region (referred to as 114 kb) and B6J littermates were generated via heterozygous-heterozygous 114 kb breeders. To avoid genetic drift, heterozygous 114 kb breeders were maintained by mating heterozygous male offspring with C57BL/6J females purchased from The Jackson Laboratory in Bar Harbor, ME, USA. Mice were SNP-genotyped using genomic DNA extracted from tail biopsies and two Taqman SNP markers: rs29383600 (50.37 Mb) and rs29459915 (50.48 Mb) (ThermoFisher Scientific, Waltham, MA, USA).

### Methamphetamine (MA)-induced locomotor activity in 114 kb congenic mice

Both female and male littermates (56-100 days old at the start of the experiment), were phenotyped for MA-induced locomotor activity. Mice were housed in same-sex groups of 2 to 4 mice per cage in standard shoebox cages and housed within ventilated racks under standard housing conditions. Colony rooms were maintained on a 12:12 h light–dark cycle. The estimated sample size required to detect a significant effect (Cohen’s d = 0.72), with 80% power (p < 0.05) was n = 25 per genotype based on the previously published phenotype in the larger congenics capturing a QTL for reduced MA-induced locomotor activity (28). Mice were tested for baseline locomotor activity on Days 1 and 2 over 60 min and then administered MA (2 mg/kg, i.p.) on Days 3, 4, and 5 and were video-recorded for distance traveled over 60 min using Anymaze (Stoelting Co., Wood Dale, IL, USA) as previously described (28). Data are presented in six, five-min bins or as the summed distance traveled over 60 min for each of the five days of injections.

### Transcriptome analysis followed by gene set enrichment analysis

Striatum punches were harvested bilaterally from 114 kb congenic mice homozygous for the congenic region (n = 8) and B6J wild-type littermates (n = 8) and processed for total RNA extractions (28). RNA samples were bioanalyzed (RIN > 8) and cDNA library was prepared using the Illumina TruSeq Standard mRNA LT (100 bp paired-end reads). The 16 samples were multiplexed and sequenced over three lanes on Illumina HiSeq 2500. The data is available on the NCBI GEO database under accession number GSE76929. FastQ files were aligned to the mouse mm10 reference genome using TopHat (33). HTSeq Python framework (34) was used to compute the read counts per gene followed by limma (35) to integrate counts and detect differentially expressed genes. For differential analysis, a linear model was used to compare gene expression between genotypes with “Cage” included as a covariate. Furthermore, between-technical replicate correlation was accounted for using the duplicateCorrelation(), limma function (36). An α level of p < 0.001 was employed. For gene set enrichment analysis of the 69 differentially expressed genes that meet the significant cutoff, Enrichr (37, 38) as used to determine the enrichment of top Kyoto Encyclopedia of Genes and Genomes **(KEGG)** pathway terms and gene ontology **(GO)** terms.

### Differential exon usage analysis followed by gene set enrichment analysis

For differential usage of exons using limma (35), reads were aligned to the Ensembl-annotated genome that contains extensive annotation of coding and non-coding exons and quantified using DEXSeq (39) with a requirement of one read count per exon bin and a minimum of at least 10 reads across all replicates. Similar to differential analysis, for each gene, a linear model was used to detect differential exon usage between genotypes with “Cage” included as a covariate and between-technical replicate correlation was accounted for using the duplicateCorrelation(), limma function (36). Differential exon usage was defined as the proportion of total normalized reads per gene that are counted within an exon bin for that gene. Statistical significance was evaluated using gene-level tests, including an F test that reflects a consensus signal across a gene and is highly powered to detect differential exon usage when more than one signal is observed across multiple exons per gene. We also report the results from a Simes multiple testing procedure that assesses all exons within a gene which is more powered to detect differential exon usage when a single strong signal is observed. An α level of p < 0.001 was employed for both the F-test and the Simes test. The 35 significant genes exhibiting differential exon usage between the 114 kb and B6J were analyzed for pathway enrichment using Enrichr (37, 38) for KEGG pathway enrichment analysis. A clustergram was used for visualization of the overlapping genes of the top enriched pathways.

### Real-time quantitative PCR (qPCR) validation

For qPCR validation of exon usage, oligo-dT primers were used to synthesize cDNA from total RNA using the same samples that were used in RNA-seq analysis using SYBR Green (ThermoFisher Scientific, Waltham, MA, USA; Cat# 4309155). The qPCR primers used for exon usage in *Hnrnph1* are shown in **Supplementary Table 1**. Each sample was run in triplicate, averaged, and normalized to its own expression level using *GAPDH* as a housekeeping gene. Differential exon usage was reported as the fold-change in 114 kb congenic mice relative to B6J littermates using the 2^-(ΔΔC^ _T_^)^ method (40).

### SDS-PAGE and Western blot

Whole striata (left and right sides) were homogenized using a hand-held homogenizer in RIPA buffer supplemented with Halt protease and phosphatase inhibitor cocktail (Thermo Fisher Scientific, Waltham, MA, USA; Cat# 78840) followed by sonication. 30 μg of protein was heated in a 70°C water bath for 10 min before loading into a 4 – 20% Criterion TGX precast Midi protein gel (Bio-Rad, Hercules, CA, USA; Cat# 5671094) for SDS-PAGE followed by wet transfer to a nitrocellulose membrane. After the transfer, all membranes were stained with ponceau S solution (0.1% ponceau S in 1% (v/v) acetic acid) for 1 minute and quickly de-stained in water to remove non-specific staining. The membranes were then imaged and densitometry analysis for total protein per lane was quantified in ImageJ2 (41). The membrane was then blocked with 5% milk for 1 h and probed with primary antibodies. hnRNP H protein expression was evaluated using three different antibodies: 1) C-term specific hnRNP H antibody (1:100,000, Bethyl, Montgomery, TX, USA; Cat #A300-511A); 2) N-term specific hnRNP H antibody (1:20,000, Santa Cruz, Dallas, TX, USA; Cat# sc-10042); and 3) hnRNP H1 antibody (1:10,000, Proteintech, Rosemont, IL, USA; Cat# 14774-1-AP). The secondary used for the Bethyl and Proteintech antibodies was donkey anti-rabbit HRP (1:10,000, Jackson ImmunoResearch Laboratories, West Grove, PA; Cat# 711-035-152) and the secondary used for the Santa Cruz one was bovine anti-goat HRP (1:10,000, Jackson ImmunoResearch Laboratories, West Grove, PA; Cat# 805-035-180). All membranes were imaged via enhanced chemiluminescence photodetection. For quantification analysis of protein expression, total protein stains were used as loading controls in normalization for immunoblotting quantification (42, 43).

### Cell culture and transfection

Human embryonic kidney (**HEK**)293T cells were grown in DMEM, high glucose with L-glutamine (Gibco, Waltham, MA, USA; Cat# 11965-092), supplemented with 10% fetal bovine serum (Gibco, Waltham, MA, USA; Cat# 26140-079), and 1% penicillin-streptomycin (Gibco, Waltham, MA, USA; Cat# 15140-122). Mouse neuro(N)2a (**N2a**) neuron-like cells were grown in 1:1 ratio of DMEM, high glucose with L-glutamine (Gibco, Waltham, MA, USA; Cat# 11965-092) to Opti-MEM (GibCO, Waltham, MA, USA; Cat# 31985-070), supplemented with 5% fetal bovine serum (Gibco, Waltham, MA, USA; Cat# 26140-079), and 1% penicillin-streptomycin (Gibco, Waltham, MA, USA; Cat# 15140-122). Cells were split every 3 to 4 days and were grown at 37°C in an atmosphere of 5% CO_2_.

For HEK293T, cells were seeded in 6-well plate at a density of 2.5 × 10^5^ cells per well 24 hours prior to transfection and would be about 90% confluent at the time of transfection. For N2A (given they grow faster than HEK293T cells), cells were seeded in 6-well plate at a density of 2 × 10^5^ cells per well. A suspension of plasmids, Lipofectamine 3000 and P3000 reagents (ThermoFisher Scientific, Waltham, MA, USA; Cat# L3000008) in Opti-MEM medium was pre-incubated at room temperature for 15 min before being added to each well of the 6-well plates. The cells were co-transfected with 1 μg of the experimental firefly luciferase construct and 0.2 μg internal pRL control plasmid expressing *Renilla* luciferase driven by the CMV promoter (Promega, Madison, WI, USA; Cat# E2261). 24 hours after transfection, HEK293T cells were detached with 0.25% of Trypsin-EDTA (ThermoFisher Scientific, Waltham, MA, USA; Cat# 25200056) and counted and re-seeded at a density of 2.5 × 10^4^ cells per well in 96-well plate. N2a cells were re-seeded at a density of 2 × 10^4^ cells per well in 96-well plate. Cells were allowed to grow for approximately 48 h before measurement of luciferase activity as described below.

### Luciferase reporter assay

We cloned the B6J *Hnrnph1* promoter sequence (Chr11: 50,375,375 to 50,378,330; mm10) starting at 2956 bp upstream of the annotated transcription start site for *Hnrnph1*. This sequence was amplified from B6J genomic DNA extracted from spleen tissue using iProof High-Fidelity PCR kit (BIO-RAD, Hercules, CA, USA; Cat# 172-5330) with the following primers containing *XhoI* and *HindIII* restriction enzyme sites in the sense and antisense primers respectively: sense (5’-GATTCTCGAGGCTCCCGTGATCAGATACAG-3’) and anti-sense (5’-GTAAAGCTTCGTCCCTTCGGTGGTCCTGGC-3’). The sequence was subsequently cloned into the multiple cloning site of pGL4.17[luc2/Neo] (Promega, Madison, WI, USA; Cat# E6721) with restriction enzymes *XhoI* and *HindIII*, placing the *Hnrnph1* promoter was inserted upstream of firefly luciferase luc2 coding sequence.

To identify variants within *Hnrnph1* between D2J and B6J mice, we used the whole genome sequence dataset (35, 36) and the online Sanger variant query tool (REL-1505 - GRCm38) to identify a total of 16 variants including three SNPs and one idel (insertion/deletion) located in the 5’ UTR of Hnrnph1 (see Table 6). To replace B6J variants with D2J variants within the 5’ UTR of *Hnrnph1*, we conducted site-directed mutagenesis using the B6J *Hnrnph1:luc2* construct as a template (Agilent QuikChange II, Santa Clara, CA, USA; Cat# 200521) to generate four luciferase reporter lines, each containing a single D2J variant. To introduce the three SNPs and the single indel all together into the 5’ UTR of *Hnrnph1*, multi-site directed mutagenesis was performed by using the *GA7546G:luc2* construct with rs221962608 SNP variant as the template (Agilent, Santa Clara, CA, USA; Cat# 200514). The mutagenic primers are provided in **Supplemental Table 2**.

**Table 1.**
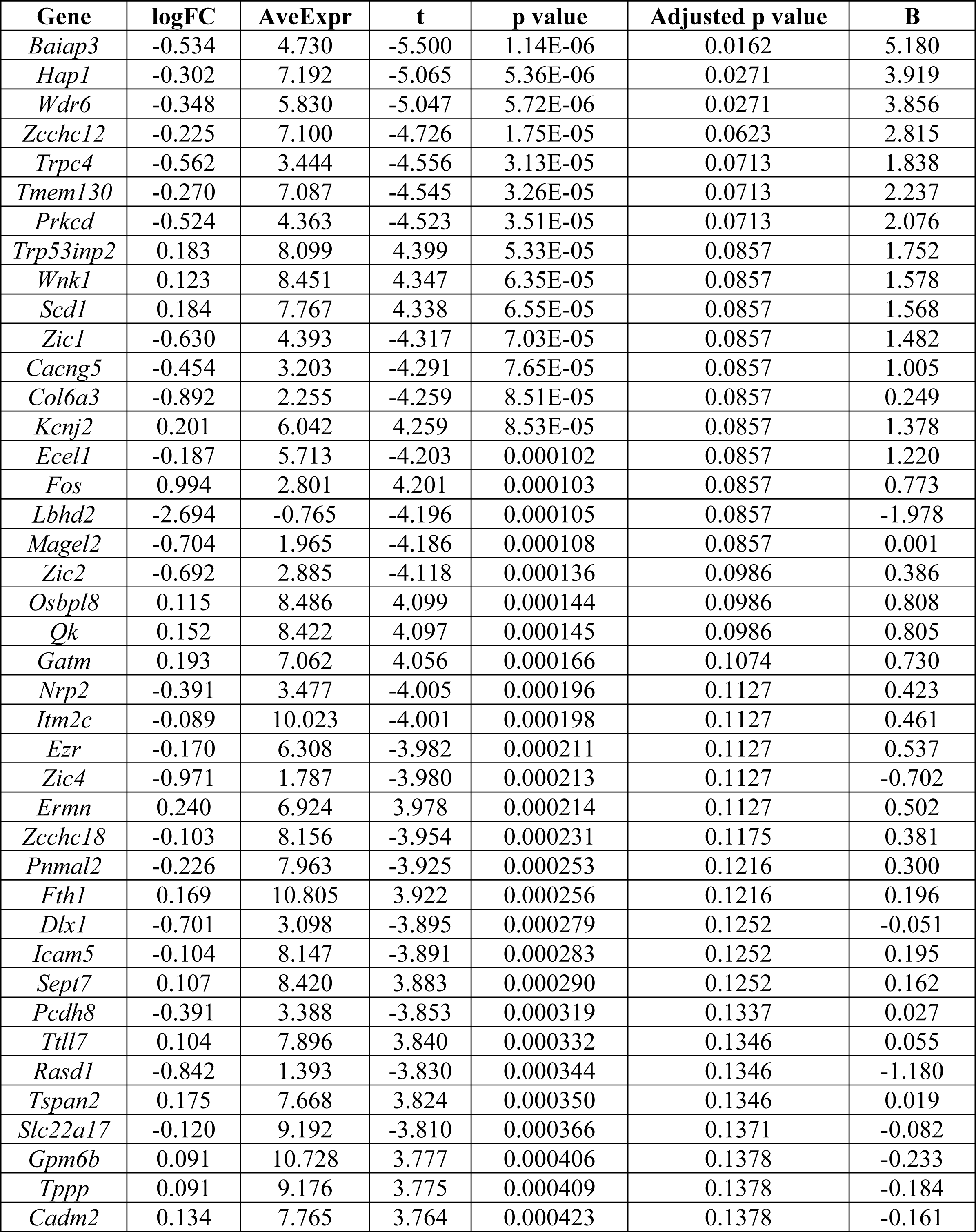
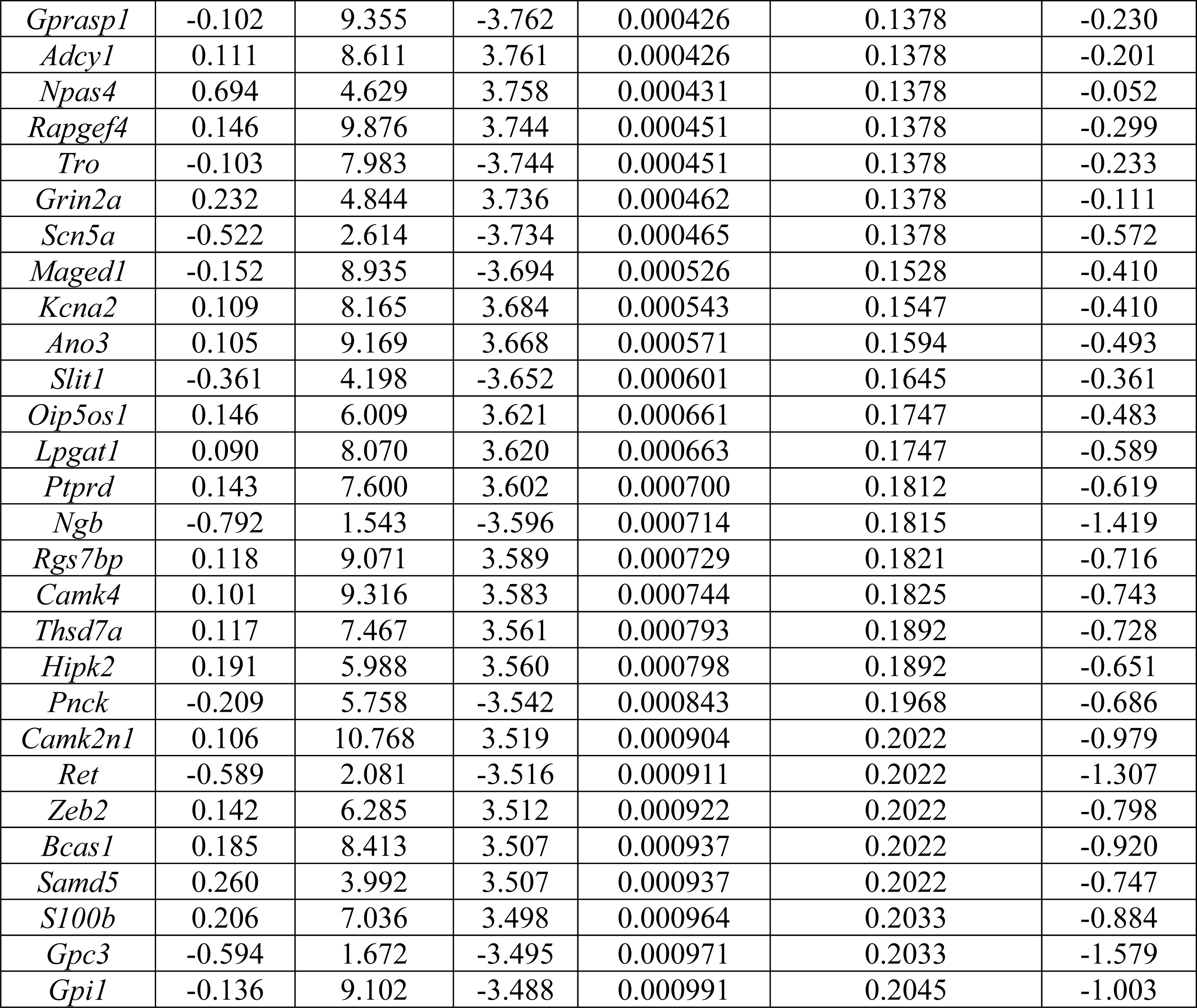
Differentially expressed genes in the striatum of 114 kb congenic mice. This table shows the 69 genes that are differentially expressed in the striatum of 114 kb congenic mice relative to the B6J wild-type littermates (p < 0.001).

**Table 2.** Pathway and gene ontology analysis of differential gene expression in 114 kb congenic mice. This table shows the top 5 KEGG pathways (**A**) and the top 5 Gene Ontology (GO) biological processes (**B**) pathways when considering the 69 differentially expressed genes (p < 0.001) between 114 kb congenic and B6J mice.

| **(A): KEGG Pathways** |  |  |  |
| --- | --- | --- | --- |
| **Name** | **p value** | **Overlap** | **Genes** |
| Oxytocin signaling pathway | 1.92E-04 | 5/154 | *Camk4, Fos, Adcy1, Kcnj2, Cacng5* |
| Circadian entrainment | 3.82E-04 | 4/99 | *Grin2a, Rasd1, Fos, Adcy1* |
| Cholinergic synapse | 6.29E-04 | 4/113 | *Camk4, Fos, Adcy1, Kcnj2* |
| cAMP signaling pathway | 8.10E-04 | 5/211 | *Grin2a, Camk4, Fos, Adcy1, Rapgef4* |
| Long-term potentiation | 1.71E-03 | 3/67 | *Grin2a, Camk4, Adcy1* |
| **(B): GO Biological Processes** |  |  |  |
| **Name** | **p value** | **Overlap** | **Genes** |
| peptidyl-threonine phosphorylation (GO:0018107) | 1.92E-04 | 4/68 | *Pnck, Wnk1, Prkcd, Hipk2* |
| regulation of non-motile cilium assembly (GO:1902855) | 3.82E-04 | 2/6 | *Sept7, Hap1* |
| peptidyl-threonine modification (GO:0018210) | 6.29E-04 | 4/89 | *Pnck, Wnk1, Prkcd, Hipk2* |
| regulation of synaptic transmission, GABAergic (GO:0032228) | 8.10E-04 | 2/11 | *Npas4, Hap1* |
| brain development (GO:0007420) | 1.71E-03 | 4/119 | *Grin2a, Zic2, Zic1, Hap1* |

**Table 3.** Genes exhibiting differential exon usage in 114 kb congenic mice. The table shows 35 genes exhibiting differential exon usage in striatal tissue from 114 kb congenic mice relative to their B6J wild-type littermates (p < 0.001). Differential exon usage was detected using either an F-test (**A**) or a Simes test (**B**) in limma (35).

| **(A): F-test ($p < 0.001$)** |  |  |  |  |  |
| --- | --- | --- | --- | --- | --- |
| **Gene** | **Fstat** | **Fstat_P** | **Fstat_FDR** | **Simes_P** | **Simes_FDR** |
| *Ppp3ca* | 2.87 | 5.12E-07 | 0.0092 | 1.30E-09 | 2.30E-05 |
| *Cdhr4, Ip6k1, Uba7, Gm20661, Gm20662* | 1.65 | 1.36E-06 | 0.012 | 0.0015 | 1 |
| *Hnrnph1* | 2.19 | 7.50E-06 | 0.045 | 0.0055 | 1 |
| *Mrps14* | 4.92 | 2.67E-05 | 0.1 | 4.10E-05 | 0.19 |
| *Pfdn5, Myg1* | 2.1 | 2.89E-05 | 0.1 | 0.028 | 1 |
| *Scn3a* | 2.14 | 4.26E-05 | 0.13 | 0.002 | 1 |
| *Chn2* | 2.53 | 5.58E-05 | 0.14 | 0.026 | 1 |
| *Myo5b* | 1.89 | 7.13E-05 | 0.16 | 0.036 | 1 |
| *Mapk9* | 2.33 | 1.20E-04 | 0.21 | 0.022 | 1 |
| *Calm2* | 3.6 | 1.30E-04 | 0.21 | 0.0024 | 1 |
| *Hnrnpm, March2* | 1.83 | 1.40E-04 | 0.21 | 0.014 | 1 |
| *Cdc42bpa* | 1.75 | 1.40E-04 | 0.21 | 0.18 | 1 |
| *Arrdc1* | 2.11 | 1.90E-04 | 0.26 | 0.21 | 1 |
| *Igbp1* | 3.98 | 3.50E-04 | 0.43 | 0.014 | 1 |
| *Sptbn1* | 1.84 | 3.60E-04 | 0.43 | 0.017 | 1 |
| *Tm4sf1* | 3.64 | 4.30E-04 | 0.49 | 0.018 | 1 |
| *Ptpn5* | 1.94 | 5.60E-04 | 0.6 | 0.2 | 1 |
| **(B): Simes ($p < 0.001$)** |  |  |  |  |  |
| **Gene** | **Fstat** | **Fstat_P** | **Fstat_FDR** | **Simes_P** | **Simes_FDR** |
| *Ppp3ca* | 2.87 | 5.12E-07 | 0.0092 | 1.30E-09 | 2.30E-05 |
| *Tspan32* | 2.96 | 0.0013 | 1 | 2.27E-05 | 0.19 |
| *Mrps14* | 4.92 | 2.67E-05 | 0.1 | 4.09E-05 | 0.19 |
| *Txn2* | 1.59 | 0.048 | 1 | 4.13E-05 | 0.19 |
| *Map2* | 1.6 | 0.0018 | 1 | 1.50E-04 | 0.53 |
| *Map4, Mtap4* | 1.57 | 0.007 | 1 | 1.80E-04 | 0.54 |
| *Slc8a3* | 2.88 | 0.0011 | 1 | 2.20E-04 | 0.57 |
| *Alpk3* | 1.76 | 0.052 | 1 | 3.80E-04 | 0.81 |
| *Jaml* | 2.74 | 0.006 | 1 | 4.10E-04 | 0.81 |
| *Dph1, Ovca2* | 1.29 | 0.15 | 1 | 4.90E-04 | 0.87 |
| *Rab21* | 2.69 | 0.15 | 1 | 5.40E-04 | 0.89 |
| *Usp24* | 0.83 | 0.86 | 1 | 9.70E-04 | 1 |

**Table 4.**
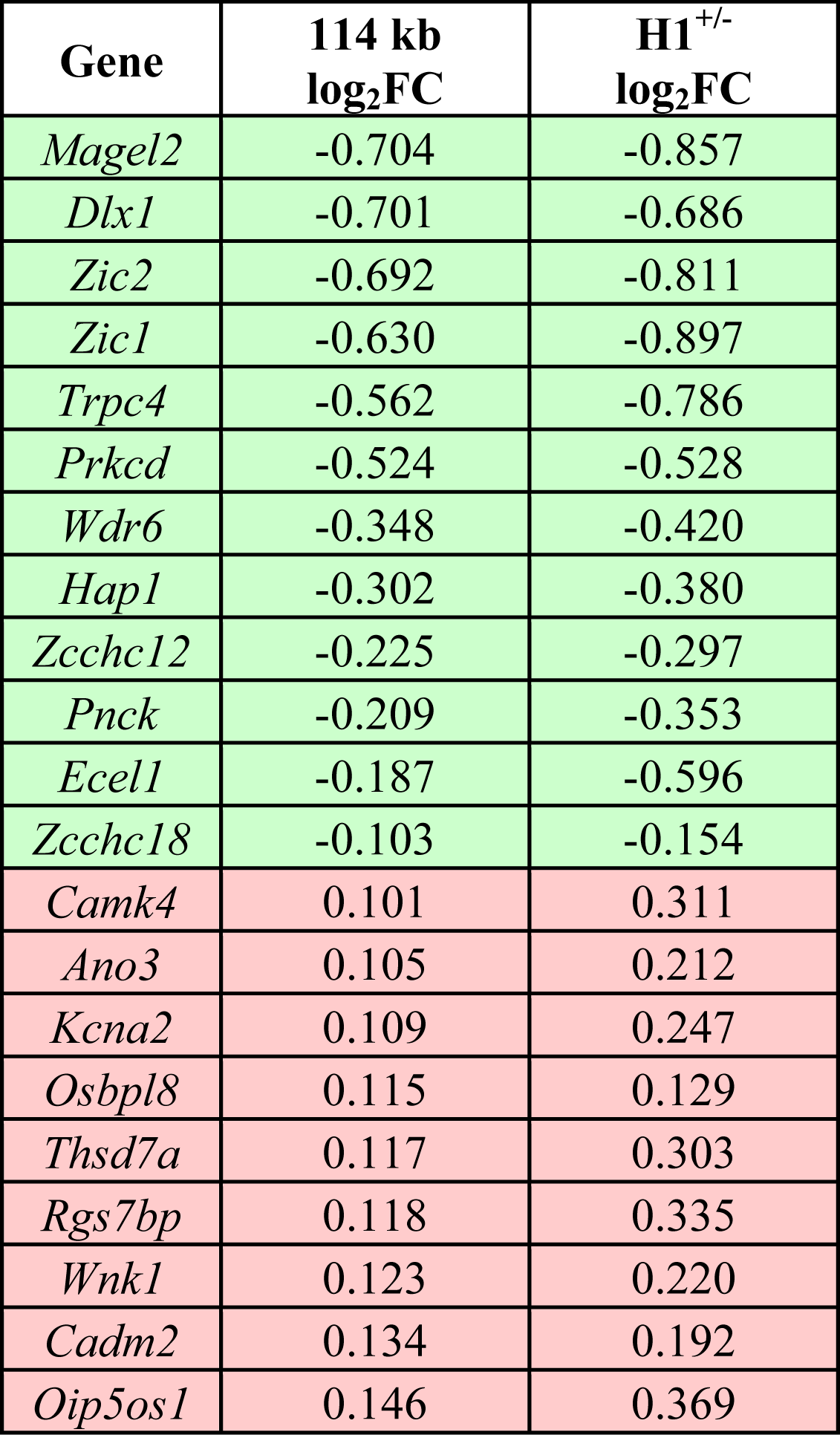
Correlation of differential expression between 114 kb congenic and H1^+/-^ mice. Table showing the log_2_FC values for the 21 overlapping genes in the 114 kb and H1^+/-^. The change in gene expression (up/down) is relative to B6J wild-type littermates.

**Table 5.**
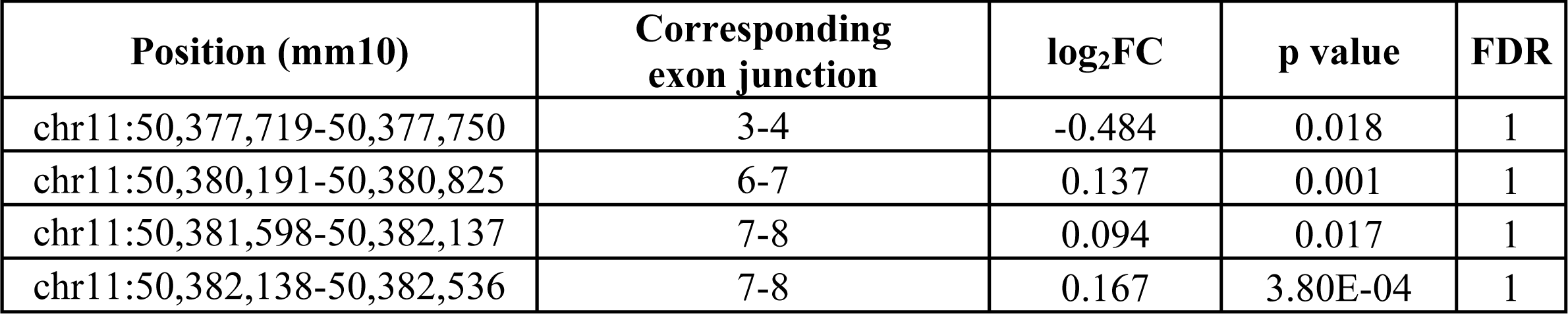
Differential exon usage of *Hnrnph1* in 114 kb congenic mice. Table showing the four exon positions of *Hnrnph1* that display differential exon usage in 114 kb congenic mice relative to B6J wild-type littermates with p < 0.05 using the Simes test. Exon usage was defined as the proportion of total normalized reads per gene that were counted within an exon bin for that gene.

**Table 6.**
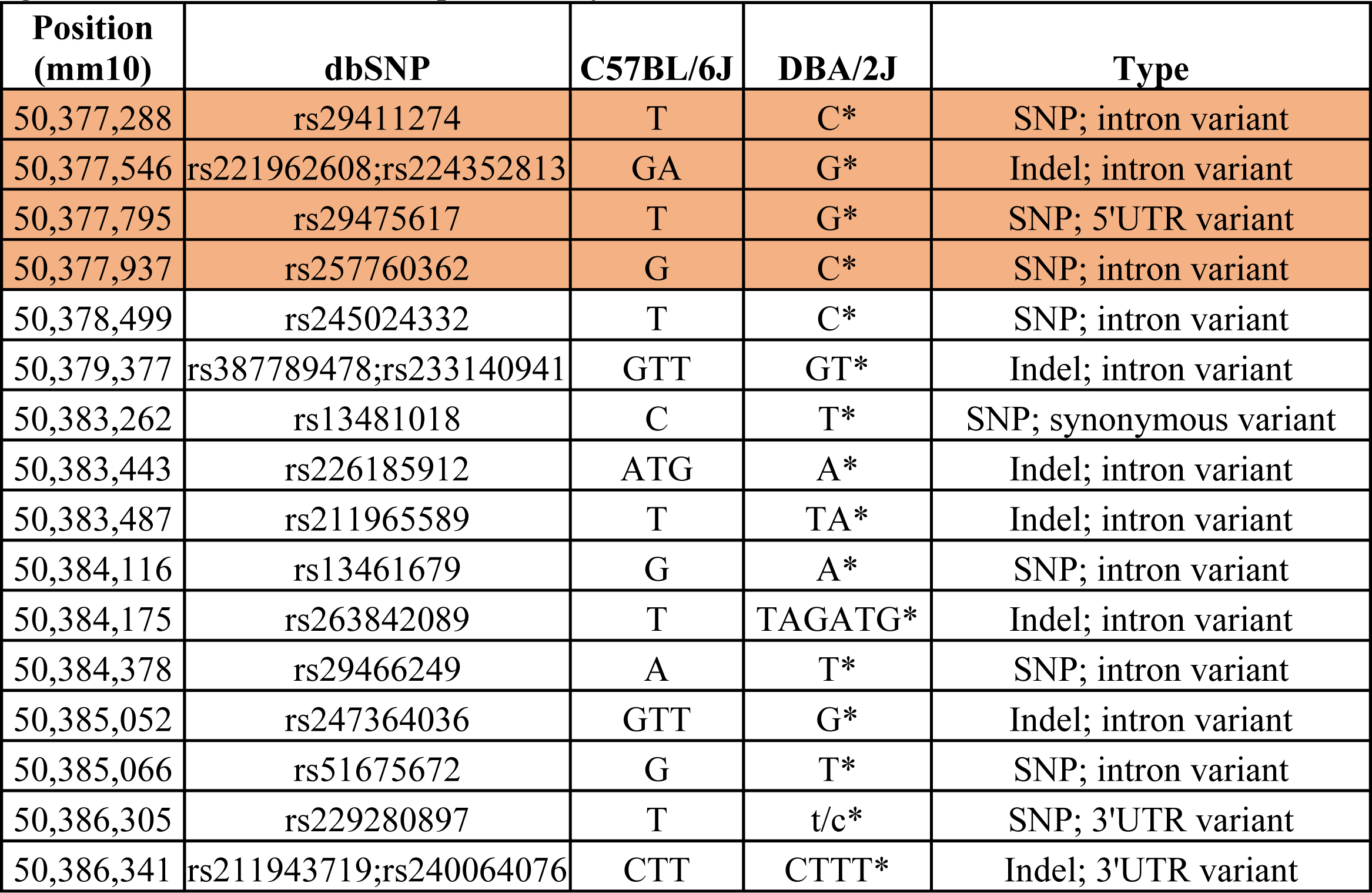
SNPs and indels in *Hnrnph1* between B6J and D2J. A query for *Hnrnph1* genetic variants (SNPs plus indels) between the B6J and D2J parental strains was conducted using the Sanger database query tool (mm10, REL-1505). The variants highlighted in orange are located in close proximity to the 5’ UTR noncoding exon 3 and were cloned and tested for functional significance in the luciferase reporter assay.

To account for background fluorescence signal, untransfected cells were used to subtract out background signal from the 96-well plate. Negative control cells were transfected with the promoter-less plasmid pGL4.17[luc2/Neo]. The Dual-Glo luciferase assay system (Promega, Madison, WI, USA; Cat# E2920) was used to measure luciferase activity. The growth medium was removed first and cells were then washed with PBS. Cell lysates were prepared by adding 50 μl of 1X passive lysis buffer to each well to lyse the cells by shaking the plate on rocker for 15 min. 20 μl of lysate from each well was transferred to wells of white opaque 96-well microplate (Corning, Corning, NY, USA; Cat# 3610). Dual-Glo LARII reagents (100 μl) were added into each well and the firefly luminescence was measured with SpectraMax i3x microplate reader after 15 min. An equal amount (100 μl) of Stop & Glo reagent was then added to the wells and Renilla luminescence was measured after 15 min.

Firefly and Renilla luciferase signal values were subtracted from the average background signal first. The background-adjusted firefly luciferase activity (FLA) for each sample was normalized to the background-adjusted internal control Renilla luciferase activity (RLA) to correct for differences in transfection efficiency and cell death. Relative luciferase activity for each experimental construct is expressed as the fold increase over the B6J *Hnrnph1:luc2* control:

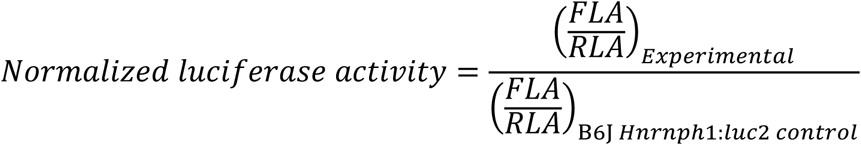

### Experimental design and statistical analyses

All data are presented as means of replicates from each experiment ± S.E.M. For experiments in which two conditions were compared, a two-tailed unpaired t-test was used to analyze the data unless otherwise specified. For the behavioral experiments in which multiple factors were evaluated, ANOVA was used and significant interactions were deconstructed along the relevant factor to examine main effects and group differences followed up with t-tests. For the qPCR validation and luciferase report assay analyses in which multiple factors were evaluated, ANOVA was used and significant interactions were deconstructed with Dunnett’s post hoc test for multiple comparisons. Statistical α level for t-test, ANOVA, and post-hoc tests were set to 0.05. The data comprising MA-induced locomotor activity were analyzed in R (http://www.R-project.org/). All other data analyses were performed in GraphPad Prism (Version 8.3.0).

## RESULTS

### Positional cloning of a 114 kb interval that is necessary for reduced MA sensitivity

In backcrossing the congenic line “Line 4a” that captured a QTL for reduced MA-induced locomotor activity and contained a heterozygous introgressed region from the D2J strain that spanned 50 Mb to 60 Mb, we monitored for recombination events at rs254771403 (50,486,998 bp, mm10). We identified a crossover event that defined 114 kb congenic region that influenced MA sensitivity. There are two protein coding genes within this 114 kb region - *Hnrnph1* and *Rufy1* **(Fig.1A)**. 114 kb congenic mice homozygous for the D2J allele within this region on an otherwise isogenic B6J background displayed no difference in saline-induced locomotor activity, compared to their wild-type homozygous B6J littermates **(Fig.1B,C)**. In response to an acute dose of MA on Day 3, 114 kb congenic mice showed reduced locomotor activity relative to B6J **(Fig.1D)**. Again, in response to a second dose of MA on Day 4, the 114 kb congenic mice showed reduced MA-induced locomotor activity compared to B6J wild-type littermates **(Fig.1E)**. No genotypic difference in MA-induced locomotor activity were observed on Day 5 after the third MA injection **(Fig.1F)** which is potentially explained by a ceiling effect on sensitization at this dose.

**Figure 1.**
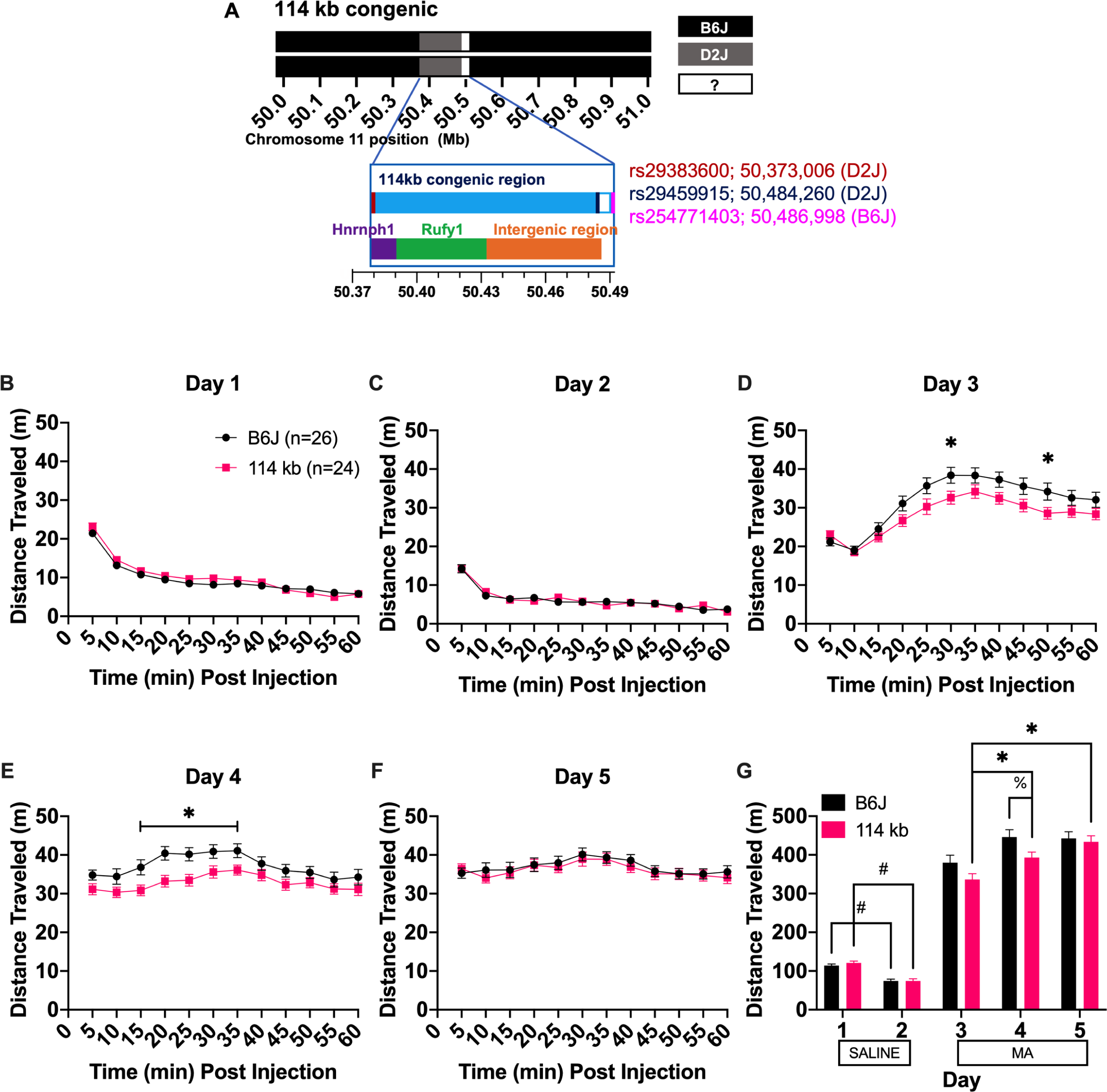
MA-induced locomotor activity in the 114 kb congenic mice. **(A):** Congenic mice containing an introgressed 114 kb region on chromosome 11 from the D2J strain on an isogenic B6J background were identified via backcrossing and screening for a recombination event at rs254771403 (50,486,998 bp; mm10) while monitoring for the retention of polymorphic rs29383600 (50,373,006 bp; mm10) and polymorphic rs29459915 (50,484,260 bp; mm10), and. Mice were then backcrossed again to B6J to generate additional heterozygotes for intercrossing (heterozygous-heterozygous breeding) which yielded heterozygous offspring as well as offspring homozygous for the 114 kb region (homozygous D2J) and wild-type homozygous B6J littermates. Behavioral analysis and results are shown for the homozygous genotypes. The 114 kb region contains two protein-coding genes: *Hnrnph1* and *Rufy1*. Mice were injected on Days 1 to 2 with saline (i.p.; panels B and C) and on Days 3, 4, and 5 with 2 mg/kg MA (panels D, E, and F). Locomotor activity was video-recorded for 60 min on each day and is presented in 5-min bins over 60 min. **(B,C):** No genotypic differences were observed in distance traveled in response to a saline injection on Day 1 [mixed ANOVA: F(1,46)_Genotype_ = 1.48, p = 0.229; F(11,506)_Genotype x Time_ = 1.48, p = 0.133] or Day 2 [mixed ANOVA: F(1,46)_Genotype_ = 0.003, p = 0.958; F(11,506)_Genotype x Time_ = 1,29, p = 0.227]. A main effect of Sex was detected for both Days 1 and 2 [F(1,46)_Day1_ = 10.52, p = 0.002; F(1,46)_Day2_ = 9.83, p = 0.003]. However, there was no significant Genotype x Sex interaction for Day 1 or Day 2 [F(1,46)_Day1_ = 0.32, p = 0.576; F(1,46)_Day2_ = 2.32, p = 0.134]. **(D):** Locomotor activity on Day 3 in response to MA is shown in 5-min bins over 60 min. 114 kb congenic mice showed reduced acute MA-induced locomotor activity relative to B6J wild-type littermates [mixed ANOVA: F(1,46)_Genotype_ = 3.41, p = 0.071; F(11,506) _Genotype x Time_ = 2.80, p = 0.002; unpaired t-test for each 5-min bin: t(48)_30min_ = 2.17, *p = 0.035; t(48)_50min_ = 2.05, *p = 0.045]. A main effect of Sex was also detected [F(1,46) = 4.32, p = 0.043], however, there was no significant Genotype x Sex interaction [F(1,46) = 2.74, p = 0.104]. **(E):** MA-induced locomotor activity on Day 4 is shown in 5-min bins over 60 min. 114 kb congenic mice showed reduced MA-induced locomotor activity relative to B6J mice [mixed ANOVA: F(1,46)_Genotype_ = 5.40, p = 0.025; F(11,506)_Genotype x Time_ = 1.96, p = 0.031; unpaired t-test for each 5-min bin: t(48)_15min_ = 2.43, *p = 0.019; t(48)_20min_ = 3.10, *p = 0.003; t(48)_25min_ = 2.90, *p = 0.006; t(48)_30min_ = 2.22, *p = 0.031]. A main effect of Sex was also detected [F(1,46) = 7.33, p = 0.009] but no significant Genotype x Sex interaction [F(1,46) = 2.05, p = 0.159]. **(F):** MA-induced locomotor activity is shown for Day 5 in 5-min bins over 60 min. 114 kb congenic mice showed no difference in MA-induced locomotor activity compared to B6J [mixed ANOVA: F(11,506)_Genotype x Time_ = 0.59, p = 0.841]. A main effect of Sex was detected [F(1,46) = 6.09, p = 0.017], however, there was no significant Genotype x Sex interaction [F(1,46) = 0.39, p = 0.535]. **(G):** Locomotor activity summed over 60 min is shown for Days 1 and 2 (saline) and on Days 3, 4, and 5 (MA). In examining habituation via changes in locomotor activity in response to saline injections on Day 1 versus Day 2, both B6J and 114 kb showed similar degree of habituation [mixed ANOVA: F(1,46)_Genotype_ = 0.47, p = 0.496; F(1,46)_Day_ = 225.51, p < 2E-16; F(1,46)_Genotype x Day_ = 1.35, p = 0.252; unpaired t-test for each day: t(50)_Day 1_ = 6.34, ^#^p = 3.26E-04; t(46)_Day 2_ = 6.64, ^#^p = 1.55E-04]. In examining sensitization of MA-induced locomotor activity across Day 3, Day 4, and Day 5, 114 kb congenic mice showed increased locomotor sensitization relative to B6J [mixed ANOVA: F(1,46)_Genotype_ = 2.74, p = 0.105; F(2,92)_Day_ = 65.71, p < 2E-16; F(2,92)_Genotype x Day_ = 5.03, p = 0.009]. Only the 114 kb congenic mice showed an increase in MA-induced locomotor activity across the three MA treatment days [Bonferroni’s multiple comparison tests (adjusted for 3 comparisons in 114 kb congenics and B6J group separately): Day 3 versus D4: t(50)_B6J_ = 2.40, p = 0.120; t(46)_114 kb_ = 2.76, *p = 0.050; Day 3 versus D5: t(50)_B6J_ = 2.45, p = 0.108; t(46)_114 kb_ = 4.52, *p = 2.57E-04; Day 4 versus D5: t(50)_B6J_ = 0.12, p = 0.999; t(46)_114 kb_ = 1.92, p = 0.368]. In addition, 114 kb congenic mice showed reduced MA-induced locomotor activity compared to B6J mice on Day 4 when compared across all five days [mixed ANOVA: F(4,184)_Genotype x Day_ = 3.96, p = 0.004; unpaired t-test for each day: t(48)_Day4_ = 2.16, ^%^p = 0.036]. A main effect of Sex was detected [F(1,46) = 9.17, p = 0.004] but there was no significant Genotype x Sex interaction [F(1,46) = 2.14, p = 0.150]. Data are represented as the mean ± S.E.M.

In examining changes in summed locomotor activity across days, both 114 kb and B6J mice showed habituation to the testing apparatus as indicated by a reduction in saline-induced locomotor activity from Day 1 to Day 2 **(Fig.1G)**. 114 kb congenic mice showed greater sensitization to repeated doses of MA from Day 3 to Day 5 **(Fig.1G)**, due to their initially lower level of acute-MA-induced locomotor activity on Day 3 compared to B6J wild-types. In comparing summed locomotor activity over 60 min across the five days, the 114 kb congenic mice showed a significant reduction in MA-induced locomotor activity compared to B6J wild-type littermates on Day 4 **(Fig.1G)**. Taken together, these results indicate that 114 kb congenic mice are less sensitive to the locomotor stimulant response to MA.

### Normalized, acute MA-induced locomotor activity and locomotor sensitization in 114 kb congenic mice

Next, we examined MA-induced locomotor activity while accounting for individual differences in baseline locomotion in response to saline on Day 2 (**Fig.2A-C**), as well as MA-induced locomotor sensitization (**Fig.2D-F**), via differences in MA-induced locomotor activity between MA treatment days. When accounting for non-drug locomotor activity, we again found that 114 kb congenic mice showed less acute MA-induced locomotor activity relative to B6J mice **(Fig.2A)**, as well as after repeated exposure to MA **(Fig.2B-C)**. After two MA injections (comparing Day 4 versus Day 3 activity), 114 kb congenic mice showed a significant decrease in locomotor sensitization early post-injection (**Fig.2D**). After three MA injections (comparing between Day 5 and Day 3), 114 kb congenic mice showed a significant increase in locomotor sensitization, starting at 20 min post-MA **(Fig.2E)**. Again, when comparing Day 5 versus Day 4, 114 kb congenic mice also showed significantly greater MA locomotor sensitization relative to B6J mice **(Fig.2F)**. The more delayed, greater sensitization in 114 kb congenic mice during the later MA injections is likely due to their initially lower MA response which takes longer to sensitize.

**Figure 2.**
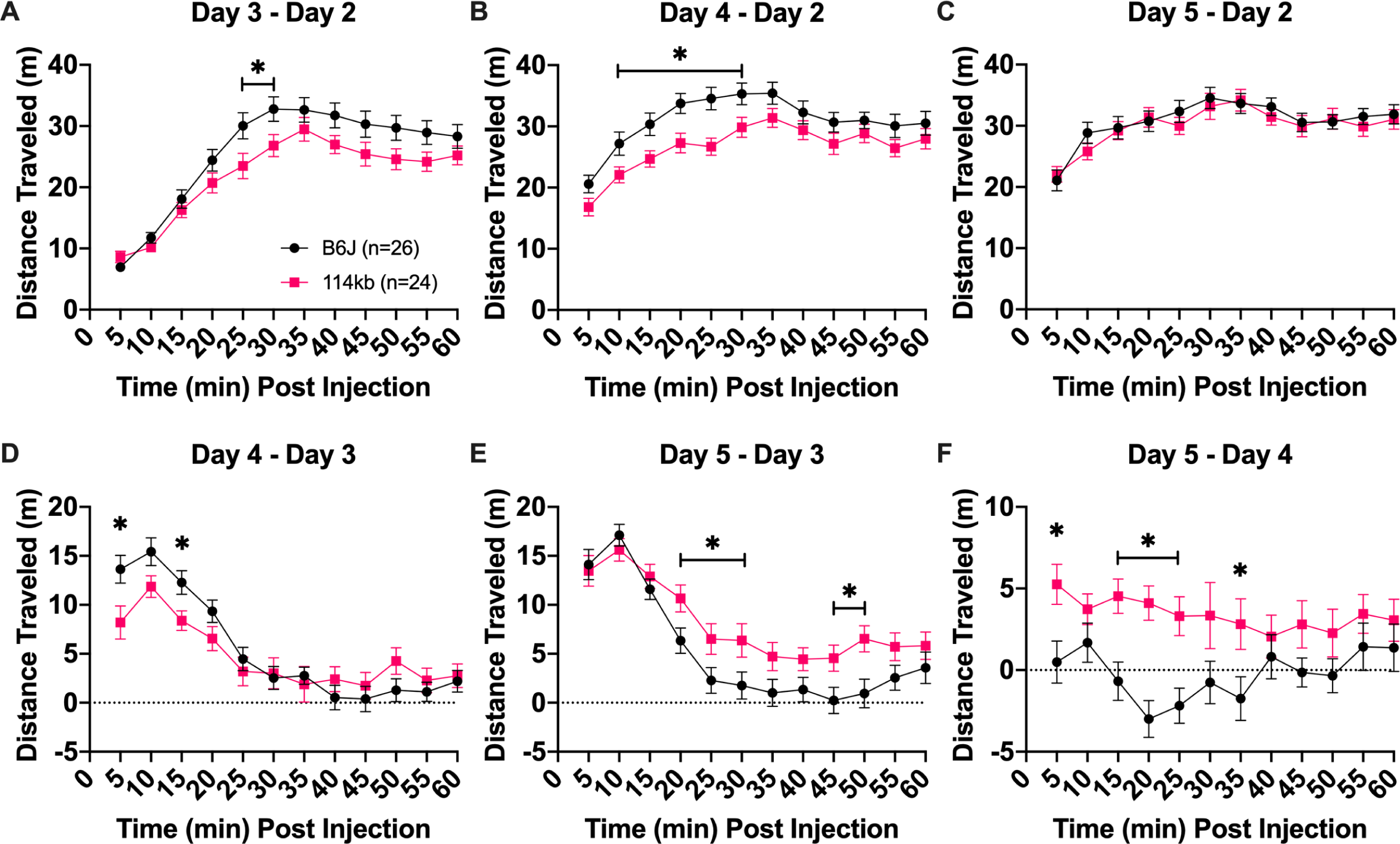
Normalized MA-induced locomotor activity and locomotor sensitization in 114 kb congenic mice. Mice were injected (i.p.) with saline on Days 1 to 2 and with 2 mg/kg MA on Days 3, 4, and 5. Locomotor activity was recorded for 60 min. **(A):** Day 3 – Day 2 distance traveled represents the acute locomotor response to MA while accounting for individual differences in non-drug, saline-induced locomotor activity. 114 kb congenic mice showed less acute MA-induced locomotor activity relative to B6J wild-type littermates [mixed ANOVA: F(1,46)_Genotype_ = 3.43, p = 0.070; F(11,506)_Genotype x Time_ = 2.36, p = 0.008; unpaired t-test for each 5-min bin: t(48)_25min_ = 2.30, *p = 0.033; t(48)_30min_ = 2.18, *p = 0.035]. **(B):** Day 4 – Day 2 distance traveled represents the locomotor response to the second, repeated dose of MA while accounting for individual differences in non-drug, saline-induced locomotor activity. 114 kb congenic mice showed less MA-induced locomotor activity relative to B6J [mixed ANOVA: F(1,46)_Genotype_ = 5.25, p = 0.027; F(11,506)_Genotype x_ Time = 1.76, p = 0.058; unpaired t-test for each 5-min bin: t(48)_10min_ = 2.15, *p = 0.037; t(48)_15min_ = 2.47, *p = 0.017; t(48)_20min_ = 2.80, *p = 0.007; t(48)_25min_ = 3.34, *p = 0.002; t(48)_30min_ = 2.24, *p = 0.030]. **(C):** Day 5 – Day 2 distance traveled represents locomotor response to the third, repeated dose of MA while accounting for individual differences in saline locomotor activity. No genotypic difference in locomotor activity was observed in response to the third dose of MA [mixed ANOVA: F(1,46)_Genotype_ = 0.17, p = 0.680; F(11,506)_Genotype x Time_ = 0.92, p = 0.523]. **(D):** Day 4 – Day 3 distance traveled is shown to represent locomotor sensitization from the first to the second MA injection. 114 kb congenic mice showed a decrease in locomotor sensitization during the first 15 min [mixed ANOVA: F(1,46)_Genotype_ = 0.47, p = 0.495; F(11,506)_Genotype x Time_ = 3.09, p = 4.92e-5; unpaired t-test for each 5-min bin: t(48)_5min_ = 2.47, *p = 0.017; t(48)_15min_ = 2.47, *p = 0.017]. **(E):** Day 5 – Day 3 distance traveled is shown to indicate the degree of sensitization observed following the third MA injection relative to the first MA exposure. 114 kb congenic mice showed an increase in locomotor sensitization relative to B6J [mixed ANOVA: F(1,46)_Genotype_ = 5.35, p = 0.025; F(11,506)_Genotype x Time_ = 1.95, p = 0.032; unpaired t-test for each 5-min bin: t(48)_20min_ = 2.30, p = *0.026; t(48)_25min_ = 2.07, *p = 0.044; t(48)_30min_ = 2.11, *p =0.040; t(48)_45min_ = 2.27, *p = 0.028; t(48)_50min_ = 2.80, p = 0.007]. **(F):** D5 – D4 distance traveled is shown to indicate which genotype continued to sensitize from the second MA injection to the third MA injection. Again, 114 kb congenic mice showed increased locomotor sensitization relative to B6J [mixed ANOVA: F(1,46)_Genotype_ = 8.50, p = 0.006; F(11,506)_Genotype x Time_ = 1.75, p = 0.061; unpaired t-test for each 5-min bin: t(48)5min = 2.67, *p = 0.010; t(48)15min = 3.28, *p = 0.002; t(48)20min = 4.59, *p = 3.17E-05; t(48)25min = 3.43, *p = 0.001; t(48)35min = 2.24, *p = 0.030]. No main effect of Sex was detected for any locomotor sensitization measures (panels A-F) [F(1,46)_Sex_ < 0.8, all p’s > 0.3]. Data are represented as the mean ± S.E.M.

### Transcriptome analysis of the striatum identifies differential usage of the *Hnrnph1* 5’ UTR in 114 kb congenic mice

To further understand the molecular mechanism of the 114 kb congenic region in reducing MA sensitivity, we performed both differential gene expression and differential exon usage analysis on RNA-seq data collected from the striatum of naïve 114 kb and their B6J wild-type littermates. We identified 69 differentially expressed genes (p < 0.001; **Table 1**). Enrichr analysis of these 69 genes for pathway and gene ontology (**GO**) terms identified enrichment for several terms potentially related to MA-induced dopamine release and behavior, including circadian entrainment, cholinergic synapse, cAMP signaling pathway, long-term potentiation, and regulation of synaptic transmission (**Table 2**).

Exon-level analysis of coding and noncoding exons (e.g., 5’ UTR noncoding exons) identified 35 genes exhibiting differential exon usage **(Table 3A,B)**, which is defined as the proportion of total normalized reads per gene that are counted within an exon bin for that gene. Enrichr analysis of the 35 genes for pathway analysis identified an enrichment of dopaminergic synapse and amphetamine addiction involving *Ppp3ca, Calm2* and *Mapk2* as well as calcium signaling and long-term potentiation involving *Ppp3ca* and *Calm2* **(Fig.3)**. Notably, *Hnrnph1* (the first protein-coding gene within the 114 kb congenic interval) was one of the top genes exhibiting differential exon usage (**Table 3A**), providing direct evidence at the exon level that one or more functional variants within *Hnrnph1* regulates exon usage and ultimately underlies differences in MA behavior. Note that *Hnrnph1* did not show a significant difference in overall gene-level transcript levels (log_2_FC = 0.016; t(14) = 0.70; p = 0.49; adjusted p = 0.99). Thus, any subsequent downstream functional changes that we report are not mediated by overall *Hnrnph1* transcript levels, but instead, by transcripts with alternative exons (either coding or noncoding; see Table 5). It should also be noted that we did not identify a significant difference in gene expression or in exon usage of *Rufy1* (the second protein-coding gene within the interval), thus limiting functional effects of D2J variants at the mRNA level to *Hnrnph1* and further supporting the candidacy of *Hnrnph1* as a quantitative trait gene underlying MA behavior (28).

**Figure 3.**
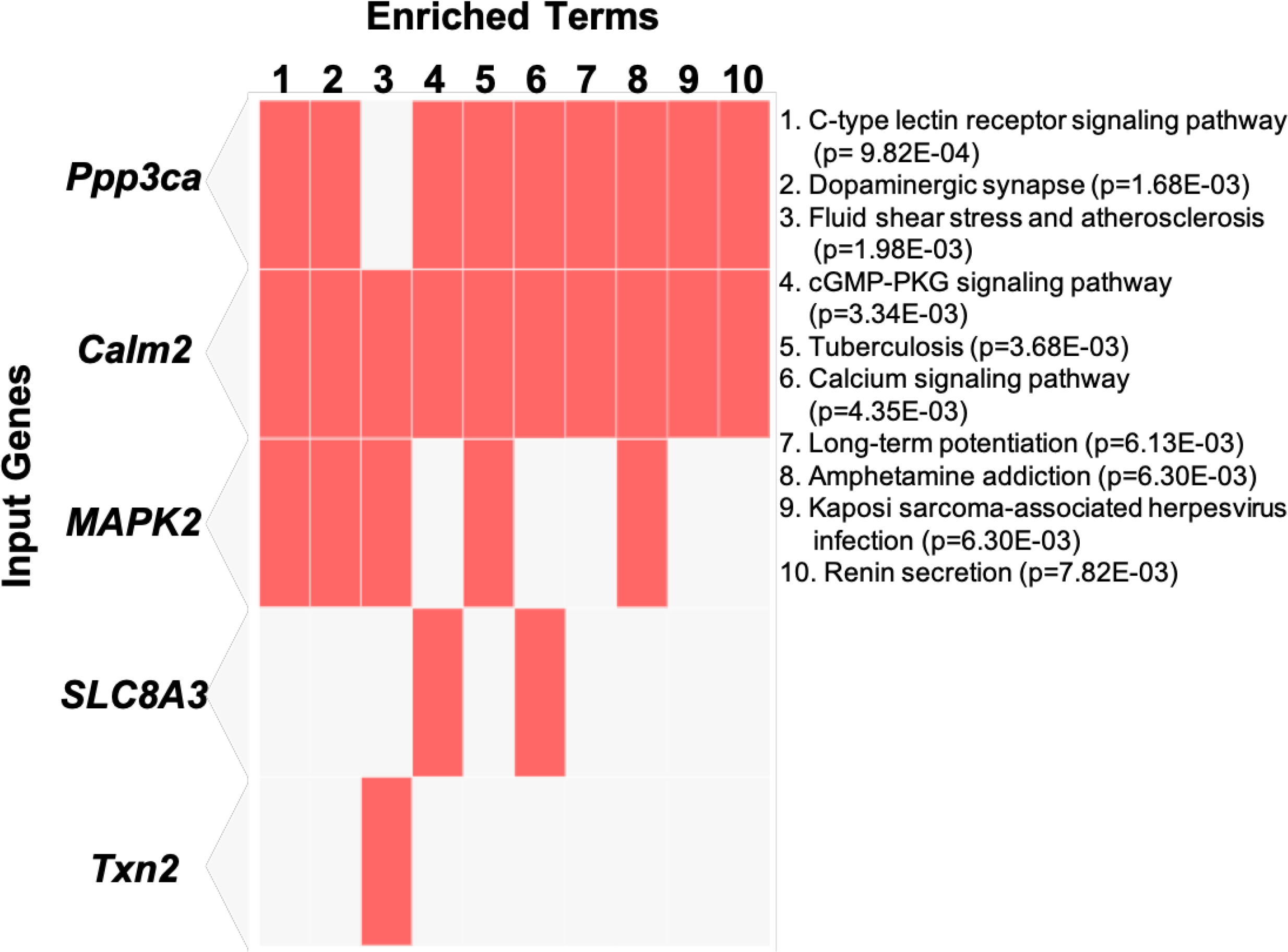
Pathway analysis of the top genes with differential exon usage. We identified 35 genes showing differential exon usage between 114 kb and B6J mice (p < 0.001). The top 10 enriched KEGG pathways along with the five genes overlapping with each of these pathways are shown in the clustergram. The enriched pathways are shown in columns and input genes are shown in the rows and cells (in orange) in the matrix to indicate if a gene is associated with a pathway.

We previously showed that mice heterozygous for a 16-bp deletion within the first coding exon of *Hnrnph1* (refer to as the H1^+/-^ mice) were less sensitive to the stimulant, rewarding, and reinforcing properties of MA and showed a reduction in MA-induced dopamine release (28, 32). To directly compare the functional effects of inheriting the deletion versus the 114 kb congenic region on gene expression and exon usage, we identified 21 overlapping genes that were differentially expressed in the striatum between 114 kb congenic mice and H1^+/-^ mice relative to their B6J wild-type littermates **(Fig.4A)**, which was significantly greater than what would be expected by chance (Fisher’s exact test: p = 9.31E-23). In addition, there were 10 overlapping genes between H1^+/-^ and 114 kb congenic mice that showed differential exon usage, relative to their B6J littermates **(Fig.4B)**. Again, this overlap was significantly greater than what would be expected by chance (Fisher’s exact test: p = 2.96E-07). Finally, in correlating differential expression for the 21 shared differentially expressed genes between 114 kb congenic mice and H1^+/-^ mice, we found a nearly perfect relationship between the magnitude and direction of change in gene expression **(Table 4 and Fig.5)**. These results suggest functionally similar effects of the 114 kb region and the H1^+/-^ allele on their overlapping, convergent transcriptome.

**Figure 4.**
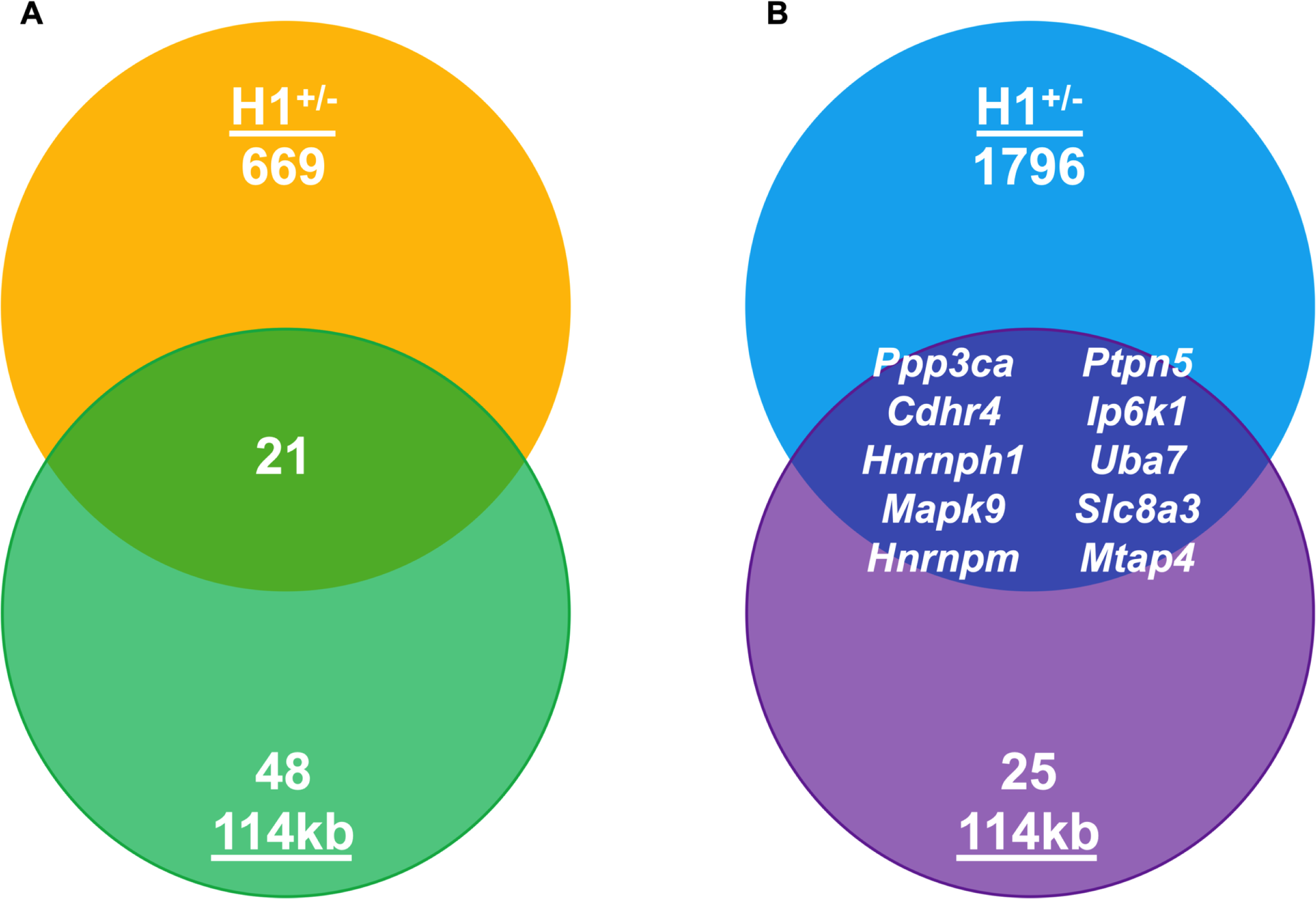
Overlap in differentially expressed genes and genes exhibiting differential exon usage between 114 kb congenic mice and H1^+/-^ mice. **(A):** Venn diagram showing the number of differentially expressed genes (**DEGs)** in 114 kb (p < 0.001) versus H1^+/-^ mice (p < 0.001). The 21 DEGs that were shared between the two are significantly greater than what would be expected by chance (Fisher’s exact test: p = 9.31E-23) as it represents nearly ½ of the 48 total DEGs identified in 114 kb congenic mice. **(B):** Venn diagram shows the number of non-overlapping and overlapping genes with differential exon usage in 114 kb congenic mice (p < 0.001) versus H1^+/-^ mice (p < 0.001). Ten genes showed differential exon usage in both the 114 kb congenic mice and the H1^+/-^ mice which is significantly greater than chance (Fisher’s exact test: p = 2.96E-07) and include *Ppp3ca, Cdhr4, Hnrnph1, Mapk9, Hnrnpm, Ptpn5, Ip6k1, Uba7, Slc8a3*, and *Mtap4*.

**Figure 5.**
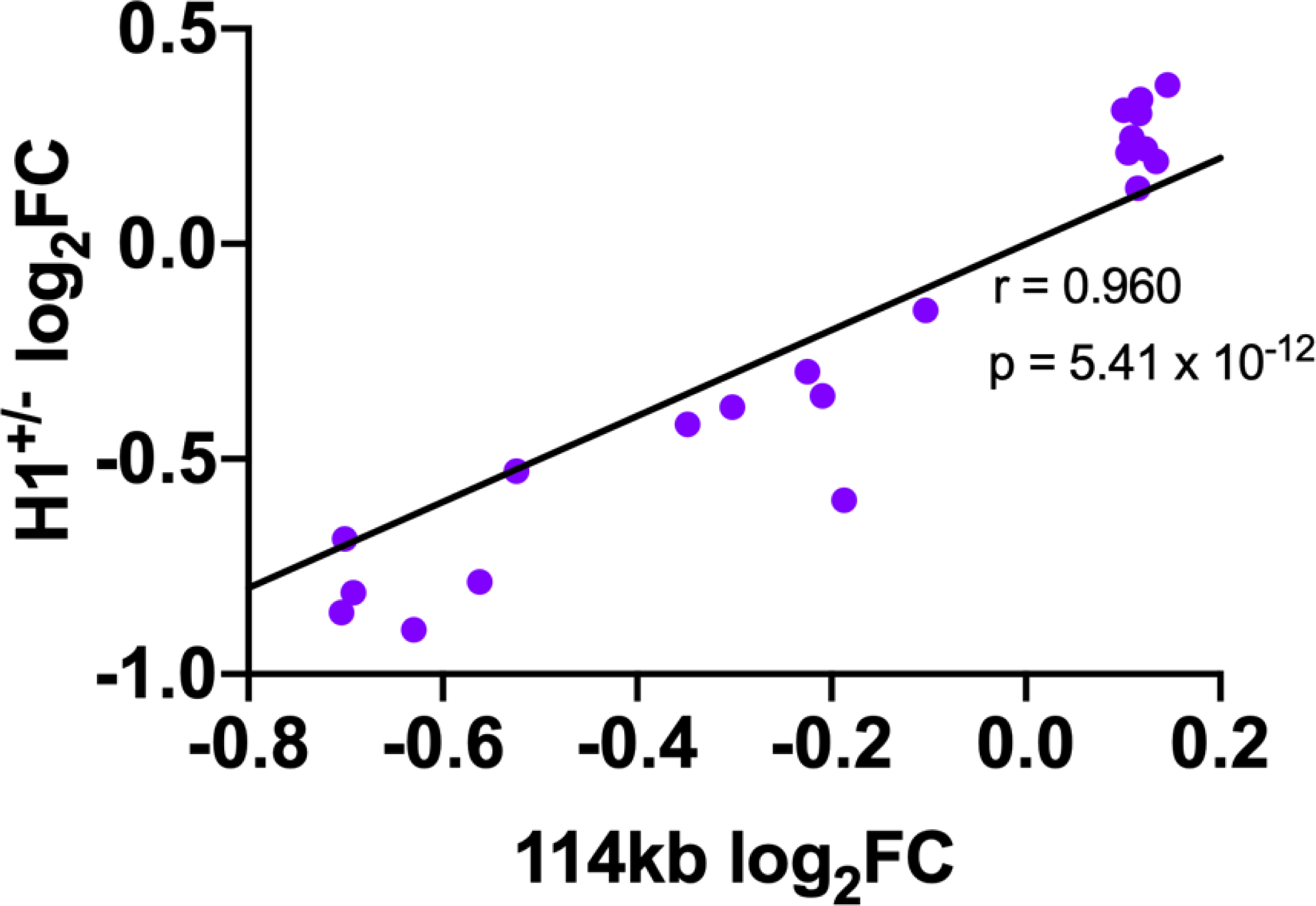
Correlation of differential expression between 114 kb congenic mice and H1^+/-^ mice. The direction and the magnitude of change in gene expression relative to B6J wildtype were nearly identical between the 114 kb congenics and H1^+/-^ mice for every gene [Pearson correlation: r = 0.960, R^2^ = 0.922, p = 5.41E-12].

The physical positions and exon numbers comprising differential exon usage of *Hnrnph1* in 114 kb relative to B6J mice are shown in **Table 5**. Note that fewer normalized reads were mapped to the junction comprising exon 3 and exon 4 in 114 kb congenic mice versus B6J (log_2_FC −0.484 = 1.4-fold decrease; **Table 5)**. Additionally, a greater number of normalized reads mapped to the junction comprising exons 6 and 7, as well as the junction comprising exons 7 and 8 in the 114 kb congenic mice relative to B6J **(Table 5)**. It is important to note that *Hnrnph2* (homolog of *Hnrnph1*) did not show differential exon usage in our analysis (F = 1.330, Fstat p = 0.198, Fstat FDR = 1 and Simes p = 0.38, Simes FDR = 0.998). To validate differential exon usage of *Hnrnph1*, we performed qPCR with primers flanking those exons exhibiting significant differential exon usage. Out of the three exon junctions, 114 kb congenic mice showed less usage of exons 3 to 4, but no difference in the other two exon junctions **(Fig.6A)**. To further validate the specific exons exhibiting differential usage, we designed primers to target exon 3 (5’ UTR noncoding exon) and exon 4 (first coding exon) separately. Less usage of the 5’ UTR noncoding exon was detected in the 114 kb congenic mice **(Fig.6B)**, with no change in the first coding exon **(Fig.6C)**. Exons 1 to 3 are noncoding exons that comprise the 5’ UTR of *Hnrnph1* (see Fig.8). Thus, we conclude that 114 kb congenic mice show a reduced number of *Hnrnph1* transcripts that include this 5’ UTR exon.

**Figure 6.**
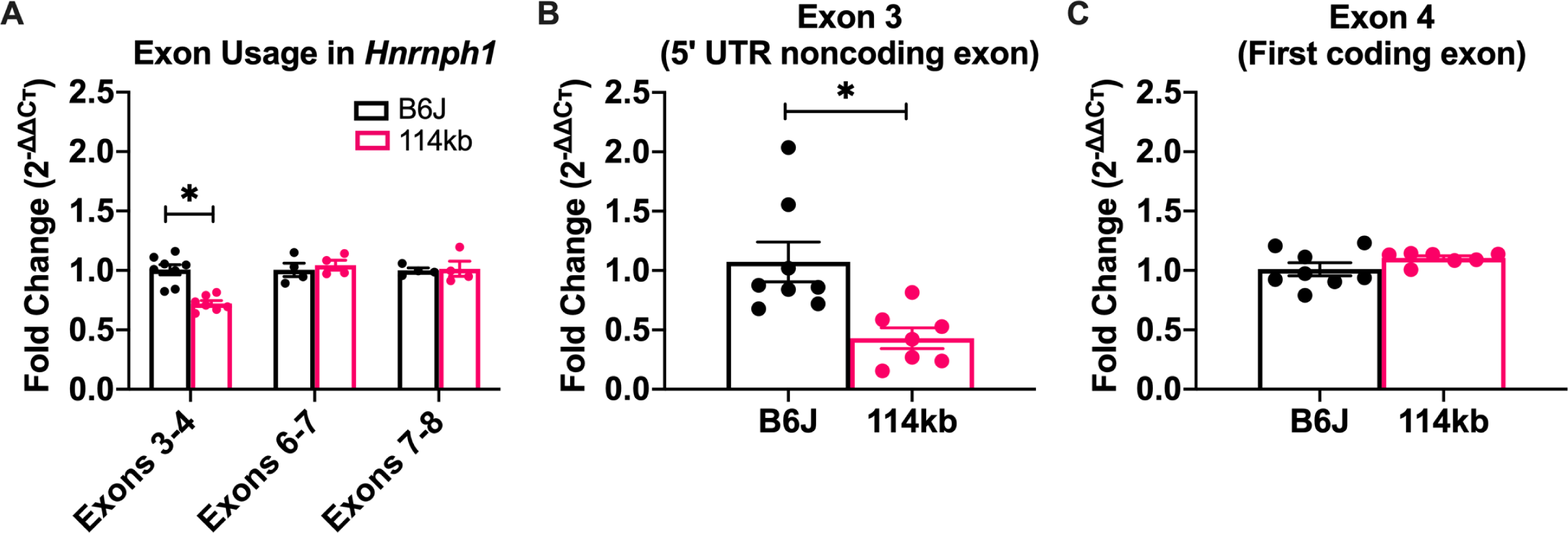
Decreased 5’ UTR noncoding exon 3 usage of *Hnrnph1* in striatal tissue of 114 kb congenic mice. *Hnrnph1* was one of the top-ranked genes showing differential exon usage as indicated via the proportion of total normalized reads counted within an exon bin between 114 kb congenic mice versus B6J wild-type littermates. To validate differential exon usage, we conducted a set of qPCR experiments targeting the exon junction or the individual exons. Primer sequences, as well as the exon junctions or individual exons they target are provided in Supplemental Table 1. **(A)** Decreased usage of the junction comprising exons 3 and 4 was detected in 114 kb congenic mice [unpaired t-test for each exon junction: t(13)_exon 3-4_ = 5.65, *p = 7.9e-5; t(6)_exon_ 6-7 = 0.58, p = 0.585; t(6)_exon 7-8_ = 0.20, p = 0.846]. **(B-C)** To further test specific exons, primers targeting either exon 3 or 4 of *Hnrnph1* were used to demonstrate that 114 kb congenic mice showed decreased usage of the 5’ UTR noncoding exon 3 [unpaired t-test: t(13) = 3.25, *p = 0.006] but no difference in the first coding exon, exon 4 [unpaired t-test: t(13) = 1.53, p = 0.150]. Data are represented as the mean ± S.E.M.

**Figure 7.**
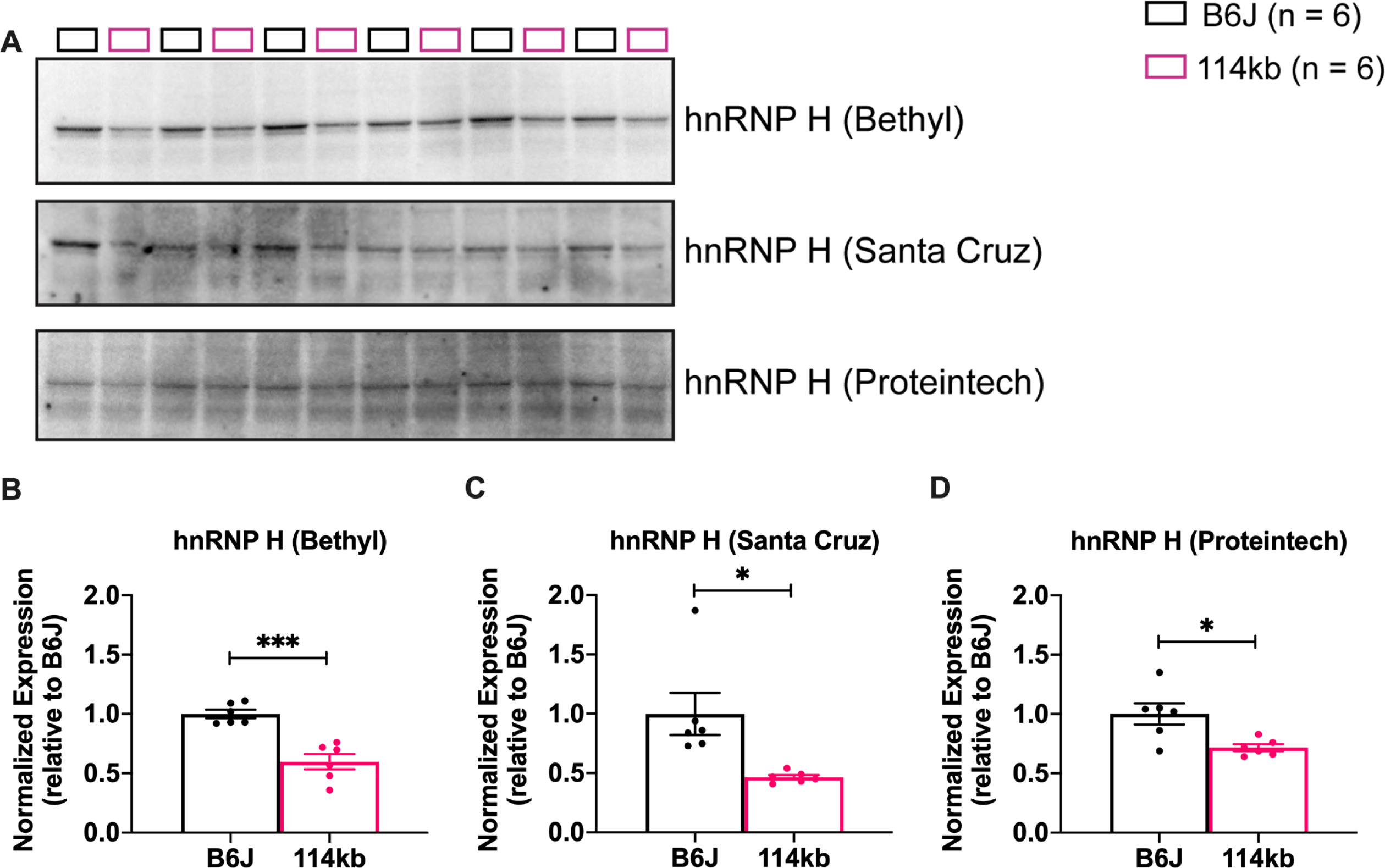
Decrease in hnRNP H protein expression in the striatum of 114 kb congenic mice. The striatum (left and right whole striatum) were dissected from the mice and protein lysates were extracted for protein quantification of hnRNP H using three anti-hnRNP H antibodies from three different companies. **(A):** Immunoblots showing hnRNP H protein expression in the striatum of 114 kb congenics and B6J mice. Three antibodies specific for hnRNP H were used. The hnRNP H antibody from Bethyl recognizes the C-terminus of the protein and the one from Santa Cruz recognizes the N-terminus. The epitope site for the antibody from Proteintech is unknown. **(B-D):** Quantification of the immunoblots shown in A showing a significant decrease in hnRNP H protein expression in the 114 kb congenic mice relative to the B6J mice when assessed with all three anti-hnRNP H antibodies [unpaired t-test: t(10)_Bethyl_ = 5.06, ***p = 0.003; t(10)_Santa Cruz_ = 2.99, *p = 0.014; t(10)_Proteintech_ = 3.02, *p = 0.013]. The expression values for hnRNP H were normalized to total protein staining by ponceau S as a loading control (See Supplemental Figure 1 for ponceau S staining images and densitometry quantification values). The normalized values were normalized to the average of B6J expression values to examine fold-change in expression relative to B6J wild-type. Data are represented as the mean ± S.EM.

**Figure 8.**
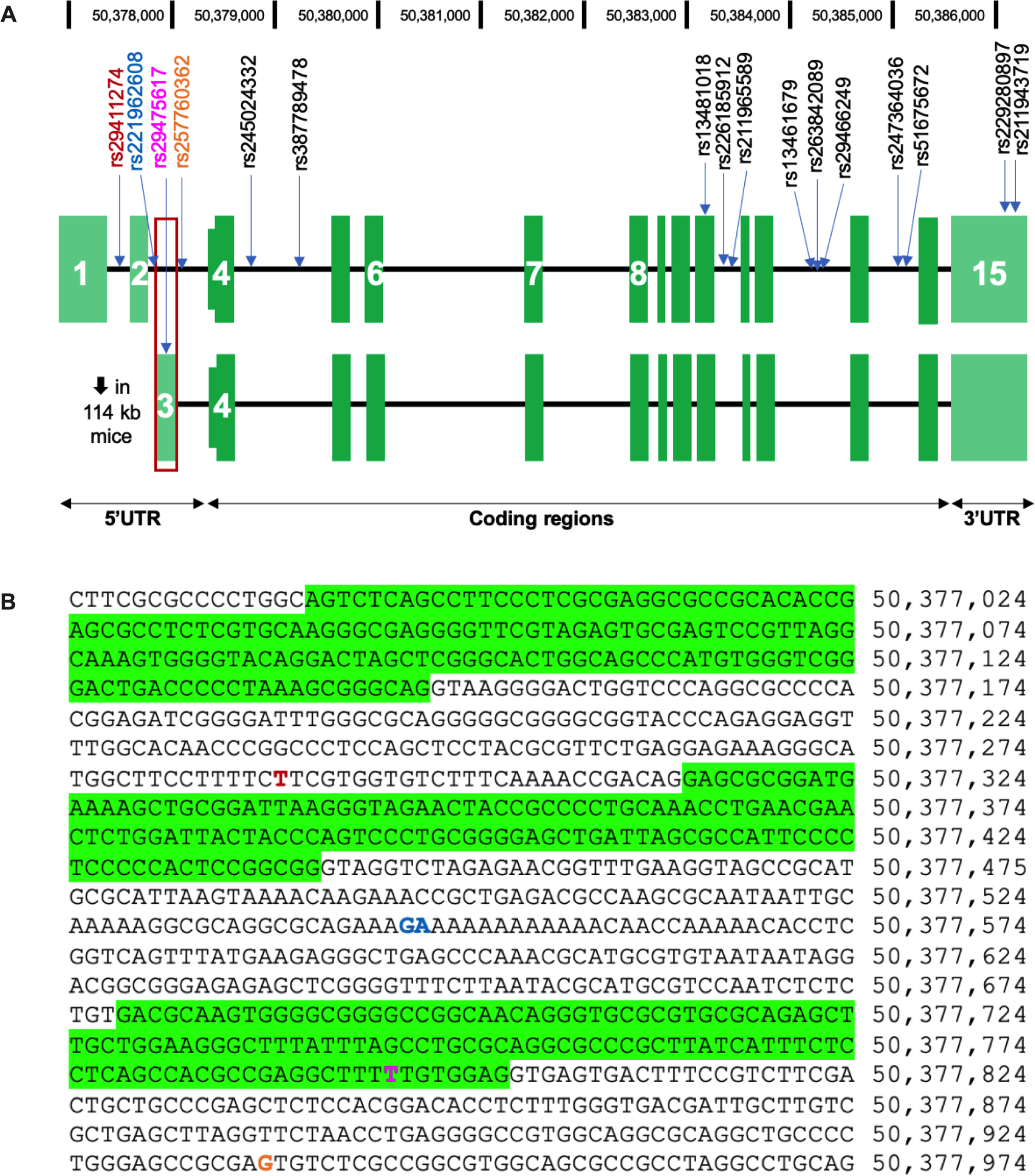
*Hnrnph1* variants between the B6J and D2J parental strains. **(A):** Schematic representation of the position of SNPs and indels in *Hnrnph1* with the exon numbers listed. The exon with decreased usage in 114 kb congenic mice is boxed in red. **(B):** Genomic sequence for part of the *Hnrnph1* 5’ UTR with positions of the three noncoding exons highlighted in green and positions of the 4 variants.

### Reduced hnRNP H protein expression in the striatum of 114 kb congenic mice

Three different antibodies for detecting and quantifying hnRNP H protein expression between 114 kb congenic mice and B6J wildtype litters were used and all samples analyzed are shown in **Fig.7A**. A significant decrease in hnRNP H immunoreactivity was detected with all three antibodies in the 114 kb congenic mice as indicated by total protein stain normalized densitometry values **(Supplemental Fig.1)** that were in turn normalized to averaged B6J values to illustrate fold-change (p = 0.003, 0.014, 0.01; **Fig.7B-D**). Thus, *Hnrnph1* variants in 114 kb congenic mice are associated with both a decrease in *Hnrnph1* 5’ UTR usage and decrease in hnRNP H protein expression.

### Identification of a set of 5’ UTR functional variants in *Hnrnph1* that decrease translation using a luciferase reporter assay

Three SNPs and one indel between D2J and B6J are located within the 5’ UTR noncoding exons and introns (**Fig.8 and Table 6)**. Given that the 5’ UTR contains a promoter element for translational regulation, we employed a luciferase reporter system to assay the strength of the *Hnrnph1* promoter in the presence of one or all four of the 5’ UTR variants in both Human Embryonic Kidney (**HEK**) 293T and Neuro2a (**N2a**) cells. We engineered a *Hnrnph1:luc2* construct by cloning 2956 bp of the B6J *Hnrnph1* promoter into the pGL4.17[luc2/Neo] promoter-less vector **(Supplemental Fig.2 and 3A; Fig.9A)**. The 2956-bp promoter increased firefly luminescence compared to the pGL4.17[luc2/Neo] vector, and this increase in signal showed a cell number dependency **(Supplemental Fig.3B)**. Using site-directed mutagenesis, we constructed 4 *Hnrnph1* promoters each possessing its own D2J variant within the 5’ UTR **(Supplemental Table 2)**. In HEK293T cells, none of the four promoters with a single D2J variant showed a significant difference between the wild-type B6J *Hnrnph1* promoter in driving luciferase expression **(Fig.9B)**. However, when all four D2J variants were introduced into the promoter, there was a significant reduction in luciferase activity compared to the B6J promoter (p = 5.68E-11; **Fig.9C)**. The findings were subsequently replicated in N2a cells (p = 1.19E-06; **Fig.9D-E)**.

**Figure 9.**
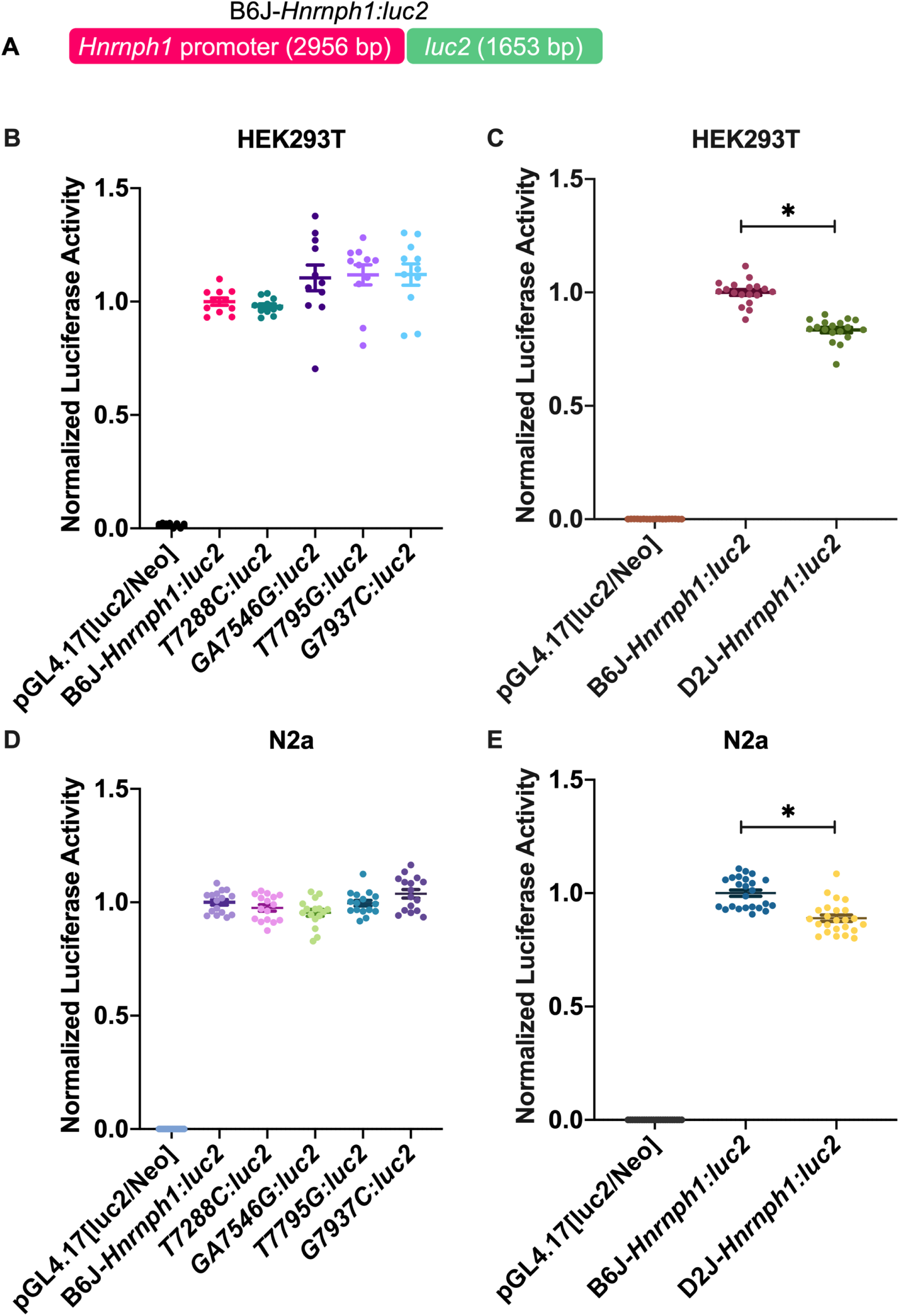
Luciferase reporter assay for the *Hnrnph1* promoter in HEK293T and N2a cells to test the functional effect of 5’ UTR variants. Firefly luciferase was used as a reporter to assess the activity of B6J-*Hnrnph1:luc2* versus promoters containing individual 5’ UTR variants from the D2J strain. *Renilla* luciferase was used as an internal control reporter. **(A):** Schematic representation of the *Hnrnph1:luc2* reporter construct that was transfected to HEK293T **(B and C)** or N2a **(D and E)** cells. **(B):** Single D2J variants were engineered within the 5’ UTR of the *Hnrnph1* promoter and had no effect on the luciferase reporter signal in HEK293T cells [one-way ANOVA comparing B6J*-Hnnrph1:luc2* against the single D2J variants: F(4,50) = 3.03, p = 0.025; Dunnett’s multiple comparison test with comparing each of the four variants with the *B6J-Hnnrph1:luc2* control group: D_Dunnett(T7288C)_ = 0.36, p = 0.988; D_Dunnett(GA7546G)_ = 1.88, p = 0.199; _DDunnett(T7795G)_ = 2.11, p = 0.125; D_Dunnett(G7937C)_ = 2.14, p = 0.117]. Data points represent the normalized luciferase signal from 2 independent replicates. **(C):** Engineering of all four D2J variants (D2J-*Hnrnph1:luc2*) in the promoter of *Hnrnph1* decreased luciferase luminescence relative to B6J-*Hnrnph1:luc2* in HEK293T cells [unpaired t-test, t(34) = 9.39, *p = 5.68E-11]. Data points represent the normalized luciferase signal from 3 independent replicates. **(D):** Singly engineered D2J variants within the promoter had no effect on luciferase reporter signal in N2a cells [one-way ANOVA comparing B6J*-Hnnrph1:luc2* against the single D2J variants: F(4,75) = 4.42, p = 0.003; Dunnett’s multiple comparison test with comparing each of the four variants with the *B6J-Hnnrph1:luc2* control group: D_Dunnett(T7288C)_ = 1.18, p = 0.584; D_Dunnett(GA7546G)_ = 2.20, p = 0.100; D_Dunnett(T7795G)_ = 0.18, p = 0.999; D_Dunnett(G7937C)_ = 1.79, p = 0.229]. Data points represent the normalized luciferase signal from 2 independent replicates. **(E):** Engineering all four D2J variants (D2J-*Hnrnph1:luc2*) within the *Hnrnph1* promoter decreased luciferase luminescence relative to the B6J-*Hnrnph1:luc2* promoter in N2a cells [unpaired t-test, t(46) = 5.59, *p = 1.19E-06]. Data points are normalized luciferase signal from 3 independent replicates. Data are represented as the mean ± S.E.M.

Taken together, these results identify a set of functional 5’ UTR variants within *Hnrnph1* that are associated with a decrease in usage of the 5’ UTR noncoding exon 3, a decrease in hnRNP H protein expression, and that reduce *Hnrnph1* 5’ UTR driven translation.

## DISCUSSION

The 114 kb region containing two protein genes (*Hnrnph1, Rufy1*) was not only necessary (28), but was also sufficient to cause a decrease in sensitivity to the locomotor stimulant response to MA **(Fig.1)**. Our prior work strongly implicated *Hnrnph1* and not *Rufy1* as the QTG responsible for the reduction in MA-induced locomotor activity (28) and we subsequently expanded the behavioral repertoire of H1^+/-^ mice to include reduced MA-induced reinforcement, reward, and dopamine release (32). Our exon-level transcriptome analysis and comparative differential gene expression analysis between the 114 kb congenics and H1^+/-^ mice further supports *Hnrnph1* as the QTG underlying the reduction in MA-induced behavior **(Fig.4 and Table 3)**. Decreased usage of the 5’ UTR noncoding exon 3 of *Hnrnph1* in 114 kb congenic mice was independently validated via qPCR **(Fig.6)**. Approximately 13% of genes in the mammalian transcriptome show differences in 5’ UTR splicing of their transcripts (44). Inheritance of the *Hnnrph1* 5’ UTR variants (along with the other *Hnrnph1* variants; **Table 6**) were associated with a decrease in striatal hnRNP H protein expression **(Fig.7)**. We identified a set of four variants within and flanking the 5’ UTR that decreased reporter expression in two different cell lines (**Fig.9**), thus identifying a set of QTVs underlying molecular regulation of *Hnrnph1* and likely MA behavior.

Identification of QTGs and QTVs is extremely rare in mammalian forward genetic studies, in particular for behavior (45), and especially when considering the identification of noncoding functional variants. In the recent identification of a noncoding functional variant underlying a *cis*-expression QTL, Mulligan and colleagues took advantage of the reduced genetic complexity of closely related C57BL/6 substrains (46, 47) to deduce and validate an intronic variant in *Gabra2* (alpha-2 subunit of the GABA-A receptor) near a splice site that drastically reduced transcript and protein expression and modulated behavior (48). Correction of the single base deletion in C57BL/6J to the wild-type C57BL/6N allele via CRISPR/Cas9-mediated gene editing restored *Gabra2* transcript and protein levels to a normal level, thus identifying the QTV (48). The vast majority of other examples that successfully identified both the QTG and the QTV involve functional coding mutations, including *Cyfip2* (49), *Grm2* (13), and *Taar1* (50). With regard to *Hnrnph1*, a human candidate gene association study identified a functional intronic variant in *OPRM1* (mRNA target of hnRNP H) that affected binding to hnRNP H to regulate splicing and was associated with the severity of heroin addiction (51).

Less than one-third of genes analyzed in human cell lines showed a direct correlation between mRNA level and protein expression (52, 53). Furthermore, *cis*-expression QTL analysis at both the mRNA and protein level in Diversity Outbred mice indicated the presence of genetic variants that regulate protein levels without affecting overall transcript levels (54). In support, our gene-level transcriptome analysis did not identify *Hnrnph1* as a differentially expressed gene; however, exon-level analysis identified decreased 5’ UTR noncoding exon usage of *Hnrnph1* in 114 kb congenic mice (**Fig.6; Table 5**) that was associated with decreased hnRNP H protein expression (**Fig.7**). Genetic variants within *cis*-regulatory elements located in the 5’ UTR and 3’ UTR of genes and *trans*-regulatory factors that bind to these elements (e.g. RNA-binding proteins (**RBPs**)) can perturb translation or overall protein abundance (52, 53, 55–57). In our study, we identified an association between decreased 5’ UTR usage and decreased hnRNP H protein expression in 114 kb congenic mice. The GC content, secondary structure, and length of the 5’ UTR can affect translational efficiency (52, 58, 58). Thus, the altered GC content within the 5’ UTR of *Hnrnph1* in 114 kb congenic mice could inhibit translation by changing the thermal stability of secondary stem loop structure (52). Because the length of the 5’ UTR determines the amount of energy needed for the ribosome to reach the AUG start site (55), a change in length of the 5’ UTR in *Hnrnph1* caused by the set of 114 kb variants could also alter protein synthesis. Also, a change in binding sites for RBPs to regulate translation could contribute to reduced hnRNP H protein in 114 kb congenic mice (59, 60). RBPs recognizes specific motifs in the 5’ UTRs, and thus, a change in the sequence of these motifs induced by these variants could alter the association of RBPs with the translation machinery at the 5’ UTR of *Hnnrph1* to control expression. The RBPDB database (61) indicates that several RBPs are predicted to bind to the 5’ UTR of *Hnrnph1*, though notably not hnRNP H itself **(Supplementary Table 3)**.

The hnRNP family of RBPs act in concert to regulate RNA metabolism and gene expression of other RBPs (62). Downregulation of hnRNP H protein in 114 kb congenic mice **(Fig.7)** could affect complexing of hnRNP H with other RBPs to regulate expression and in turn, alter recruitment of RBPs involved in splicing of the 5’ UTR of *Hnrnph1*. As an example, hnRNP A1 is predicted to bind to the 5’ UTR of *Hnrnph1* (61) and this RBP is known to cooperate with hnRNP H in mediating gene splicing (63). Thus, although *Hnrnph1* is not predicted to be a direct target of hnRNP H, reduced hnRNP H protein could alter the expression of and coordination with other RBPs to regulate 5’ UTR usage of *Hnrnph1*.

To summarize, we provide further causal behavioral and molecular evidence for *Hnrnph1* as a QTG for MA sensitivity by demonstrating that inheritance of a small, 114 kb chromosomal region was not only *necessary* (28) but also *sufficient* to induce the behavioral phenotype. We provide functional evidence for a set of four QTVs within the 5’ UTR that could plausibly reduce 5’ UTR usage and the amount of hnRNP H protein; however, demonstrating that these four variants are sufficient alone to induce these functional effects on *Hnrnph1* molecular regulation and behavior would require the generation of a mouse model containing only these four variants. The relatively subtle effect of the 114 kb region on behavior raises the question of whether modeling these variants *in vivo* is worth the effort – convergent evidence provides strong evidence that we have identified a set of QTVs underlying functional molecular and behavioral changes. Future studies involving cell type-specific knockdown of *Hnrnph1* using the Cre-loxP system will efficiently and more selectively model the effects of reduced hnRNP H protein expression on MA-induced behavior.

## NONSTANDARD ABBREVIATIONS

(PUD): psychostimulant use disorder
(MA): methamphetamine
(QTL): quantitative trait locus
(SUD): substance use disorders
(D2J): DBA/2J
(B6J): C57BL/6J
(H1^+/-^): *Hnrnph1* mutants
(QTV): quantitative trait variant
(qPCR): real-time quantitative PCR

## AUTHOR CONTRIBUTIONS

Q. T. Ruan, N. Yazdani, K.K. Szumlinski, P. E. A. Ash, B. Wolozin, and C. D. Bryant designed research; Q. T. Ruan, N. Yazdani, and E. R. Reed analyzed data; Q. T. Ruan, N. Yazdani, J. A. Beierle, L. P. Peterson, and K. P. Luttik performed research; P. E. A. Ash and B. Wolozin contributed new reagents or analytical tools; Q.T. Ruan and C. D. Bryant wrote the paper.

## Supplemental Data

**Supplemental Figure 1.**
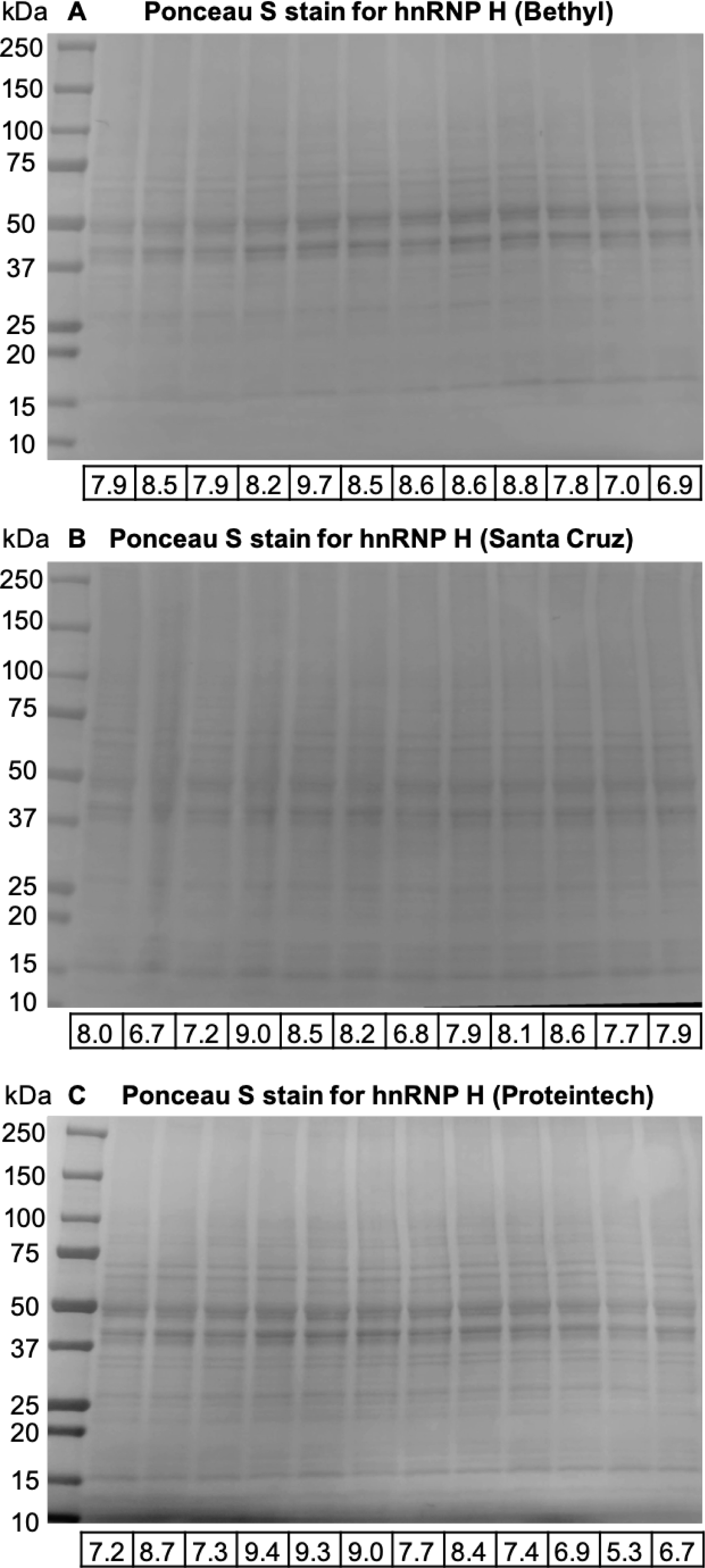
Ponceau S staining for total protein normalization used in immunoblot quantification. Total protein stains via ponceau S instead of a housekeeping gene were used as loading controls. The densitometry value for the total protein stains in each lane is indicated below each immunoblot. These are the values that were used for normalization. **(A-C):** Ponceau S total protein staining for quantification hnRNP H protein expression.

**Supplemental Figure 2.**
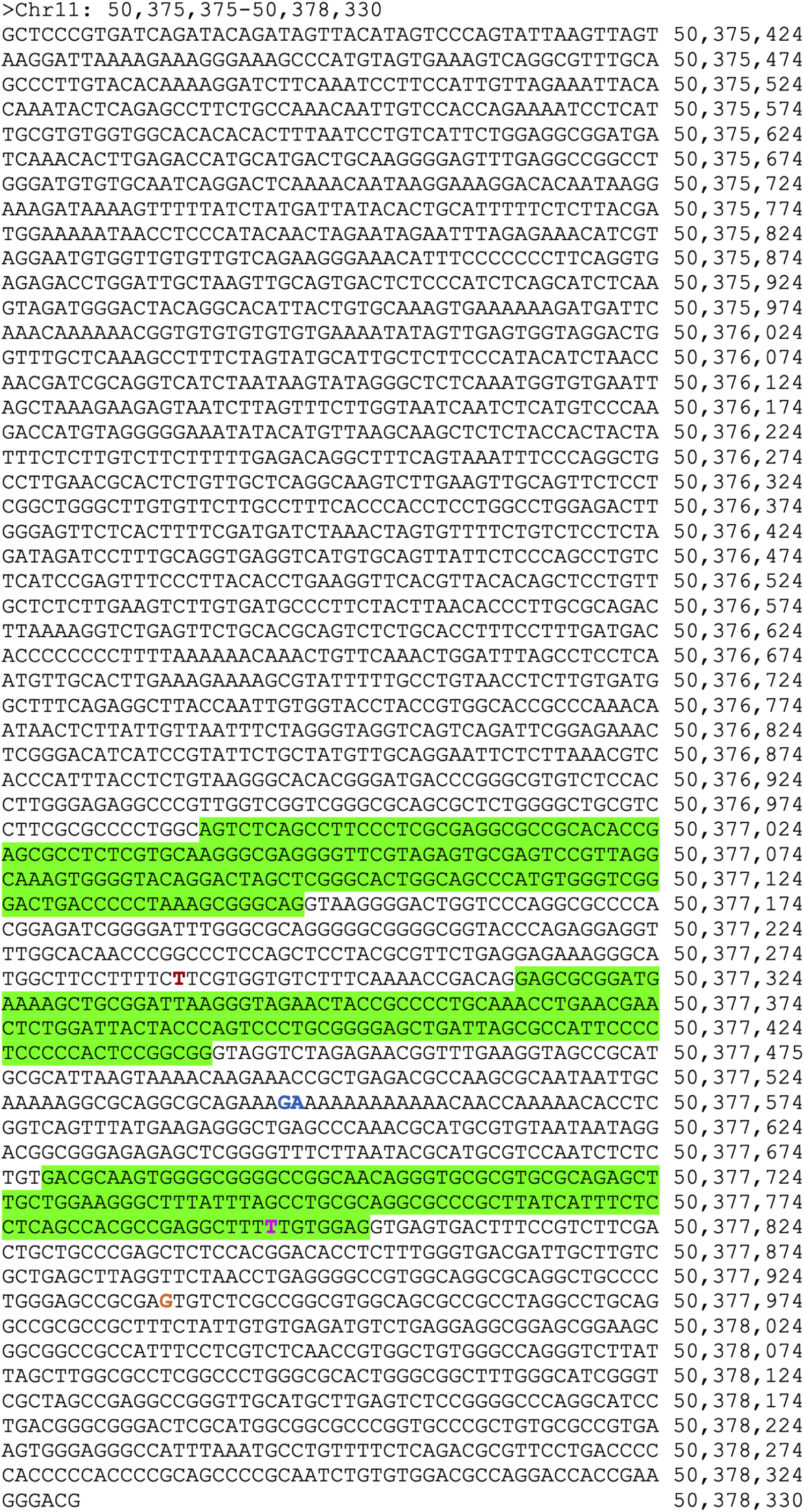
DNA sequence of the cloned *Hnrnph1* promoter. The promoter contains the DNA sequence 2956 bp upstream of transcription start site which corresponds to chromosome 11 nucleotide position 50,375,375 to 50,378,330 (mm10). The nucleotides indicated in different colors represent the positions of the SNPs, and the nucleotides indicated in blue represent the positions of the indels. The exons are highlighted in green.

**Supplemental Figure 3.**
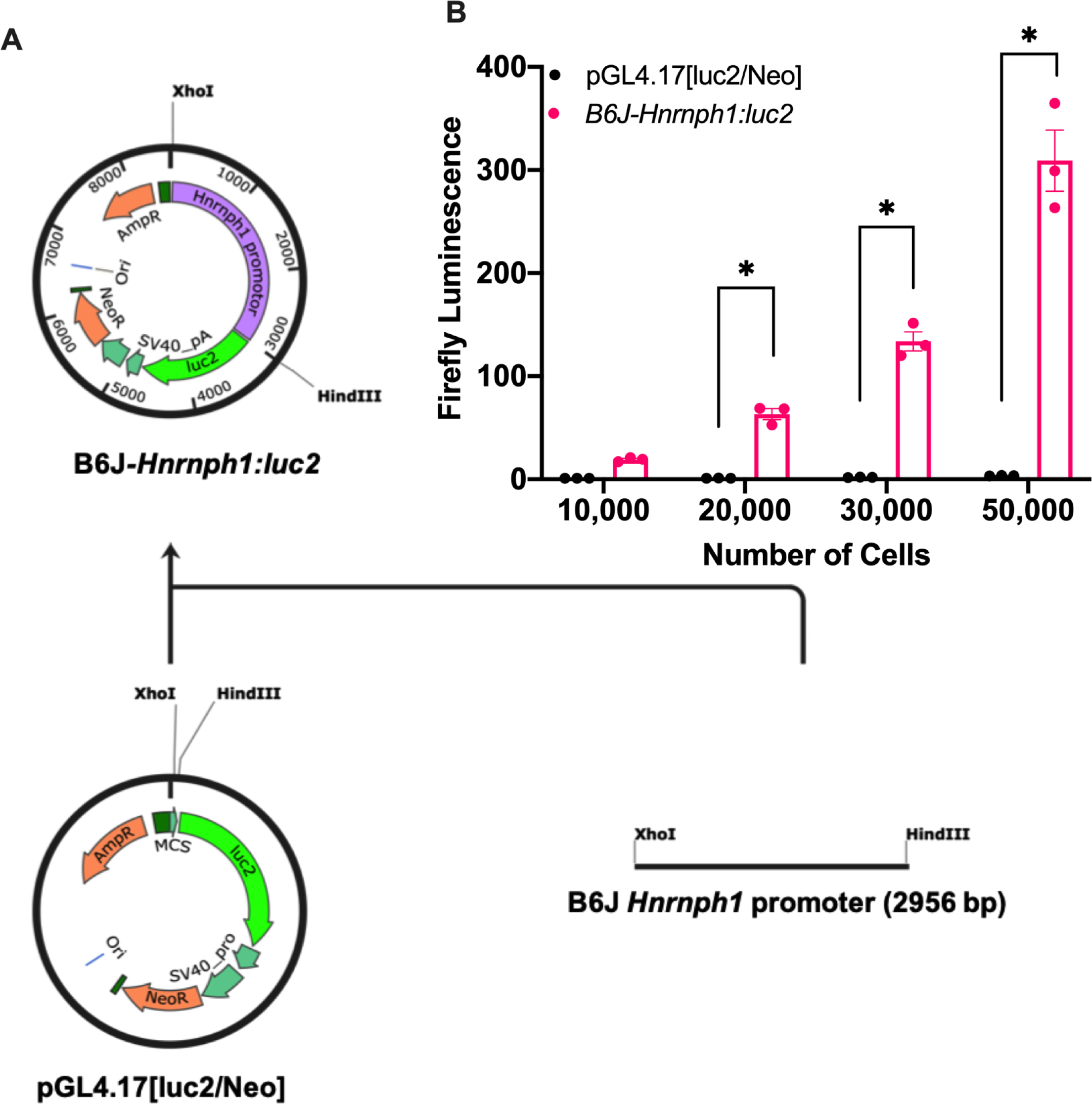
*Hnrnph1:luc2* reporter assay in HEK293T cells. **(A):** Schematic representation of the procedure for cloning the *Hnrnph1* promoter into pGL4.17[luc2/Neo]. The promoter region of *Hnrnph1* was PCR-amplified followed by restriction enzyme digest by *XhoI* and *HindIII* for insertion into pGL4.17[luc2/Neo]. *The Hnrnph1* promoter was defined as 2956 bp upstream of the transcriptional start site of *Hnrnph1* and was fused to the firefly luciferase (luc2) to make the *Hnrnph1:luc2* reporter. **(B):** Different numbers of HEK293T cells were seeded to determine whether the cloned *Hnrnph1* promoter could drive expression of firefly luciferase, *luc2*. Firefly luminescence increased as the cell number increased [two-way ANOVA: F(3,16) = 64.19, p = 3.86E-09; Bonferroni’s multiple comparison test: t(16)_10,000_ = 1.15, p > 1; t(16)_20,000_ = 2.94, *p = 0.005; t(16)_30,000_ = 8.35, *p = 1.25e-6; t(16)_50,000_ = 19.36, *p = 6.30e-12]. Data are represented as the mean ± S.E.M.

**Supplemental Table 1.**
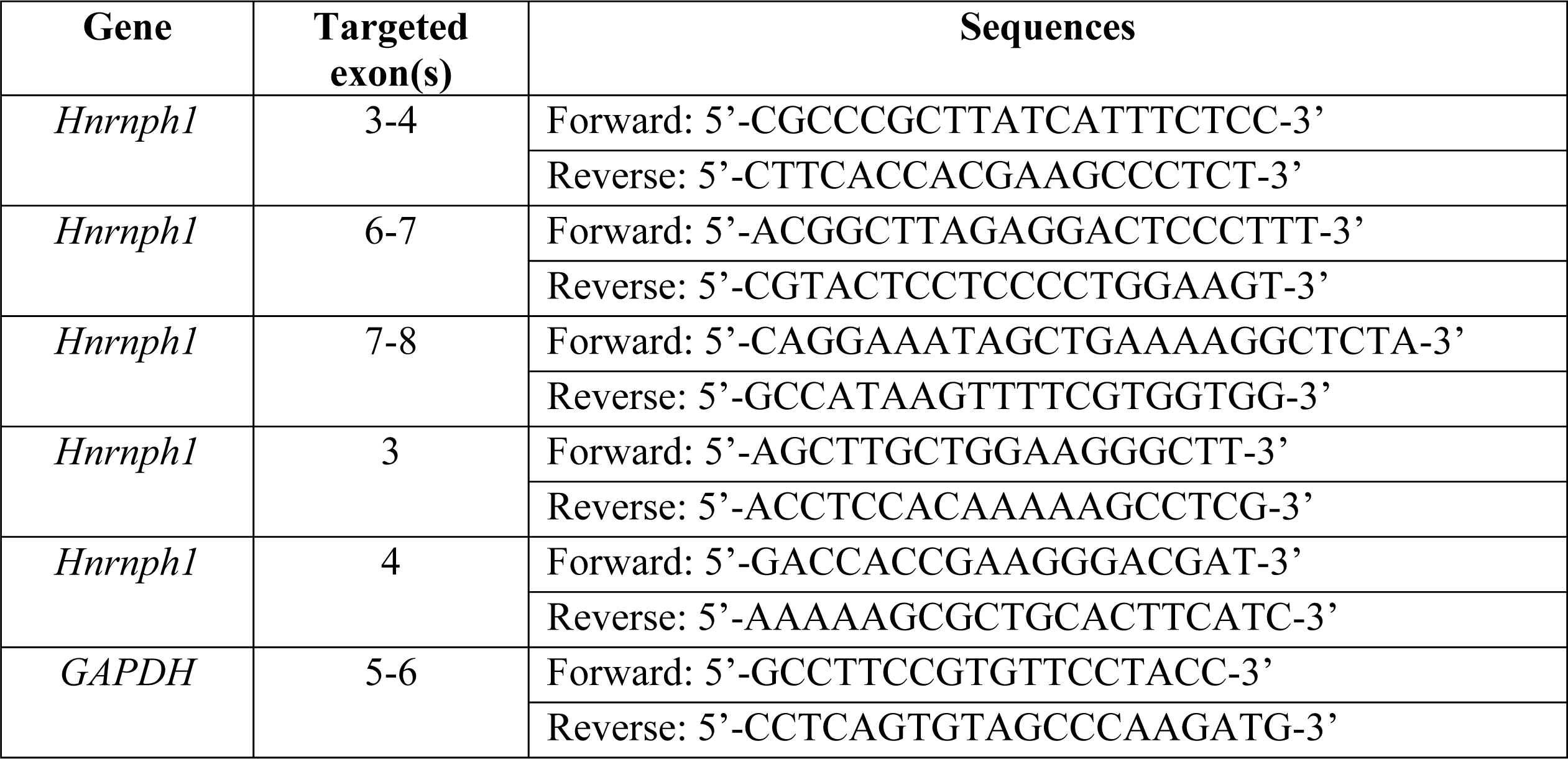
RT-qPCR primers for exon usage.

**Supplemental Table 2.**
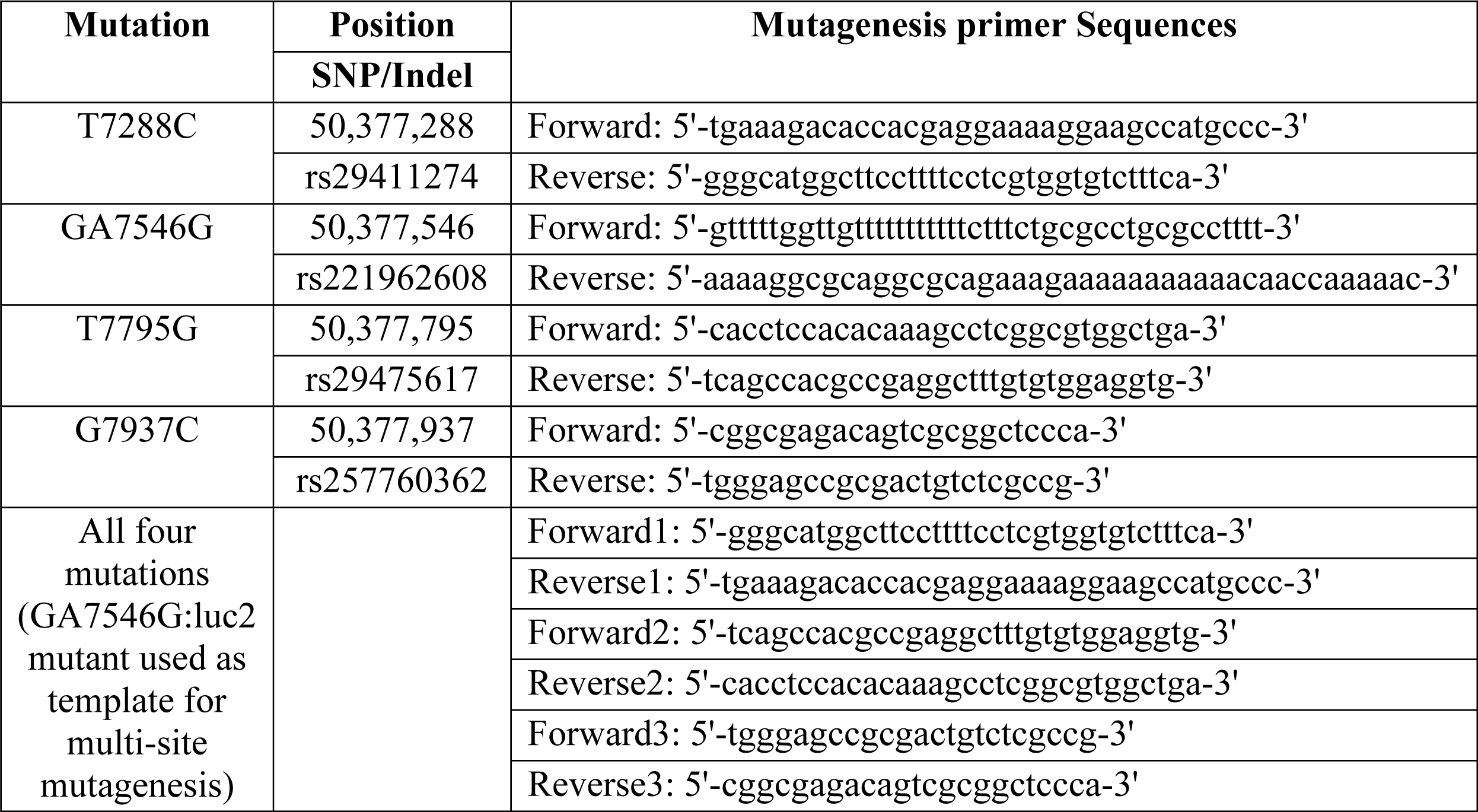
Primers used for site directed mutagenesis.

**Supplemental Table 3.**
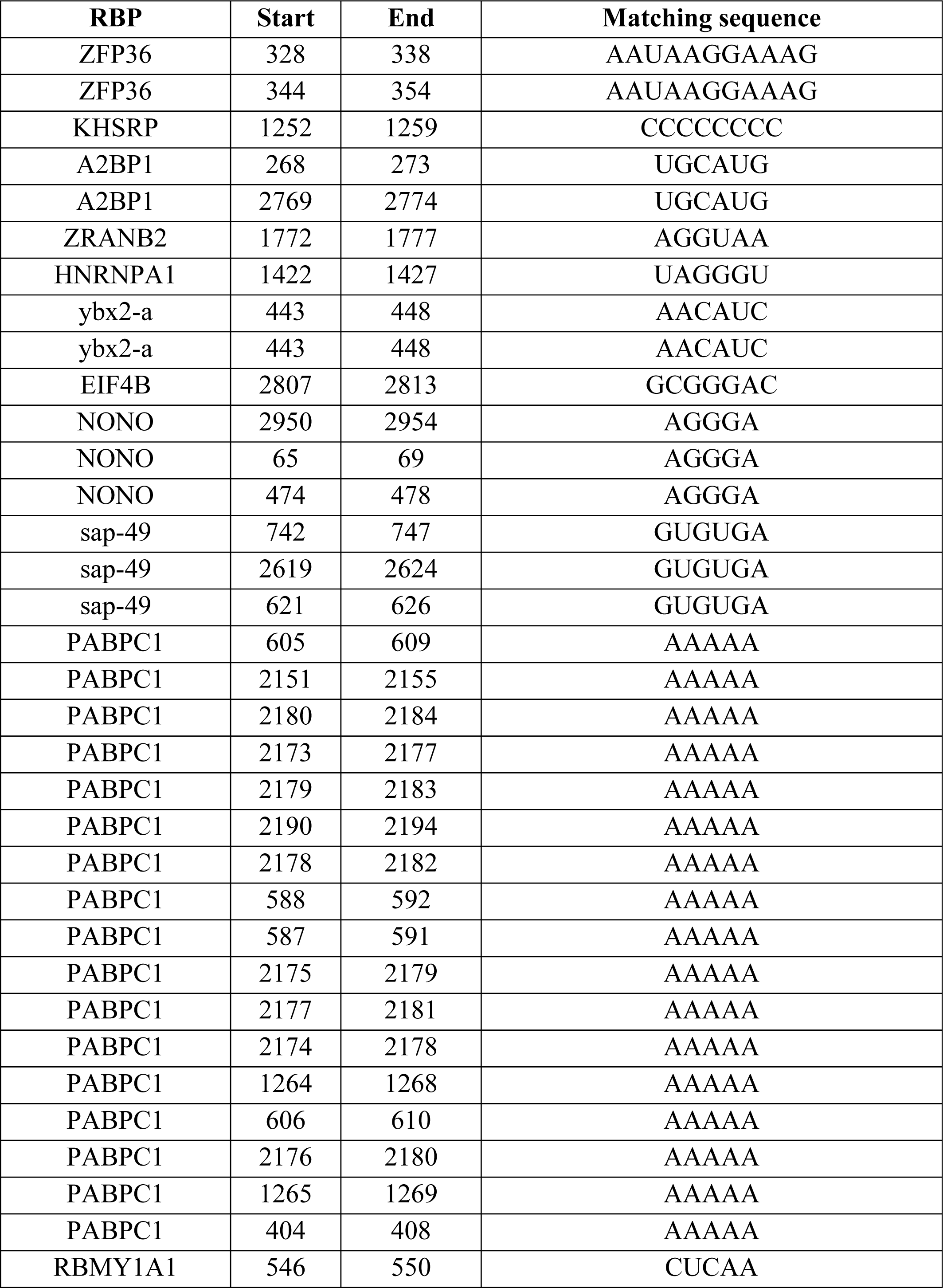
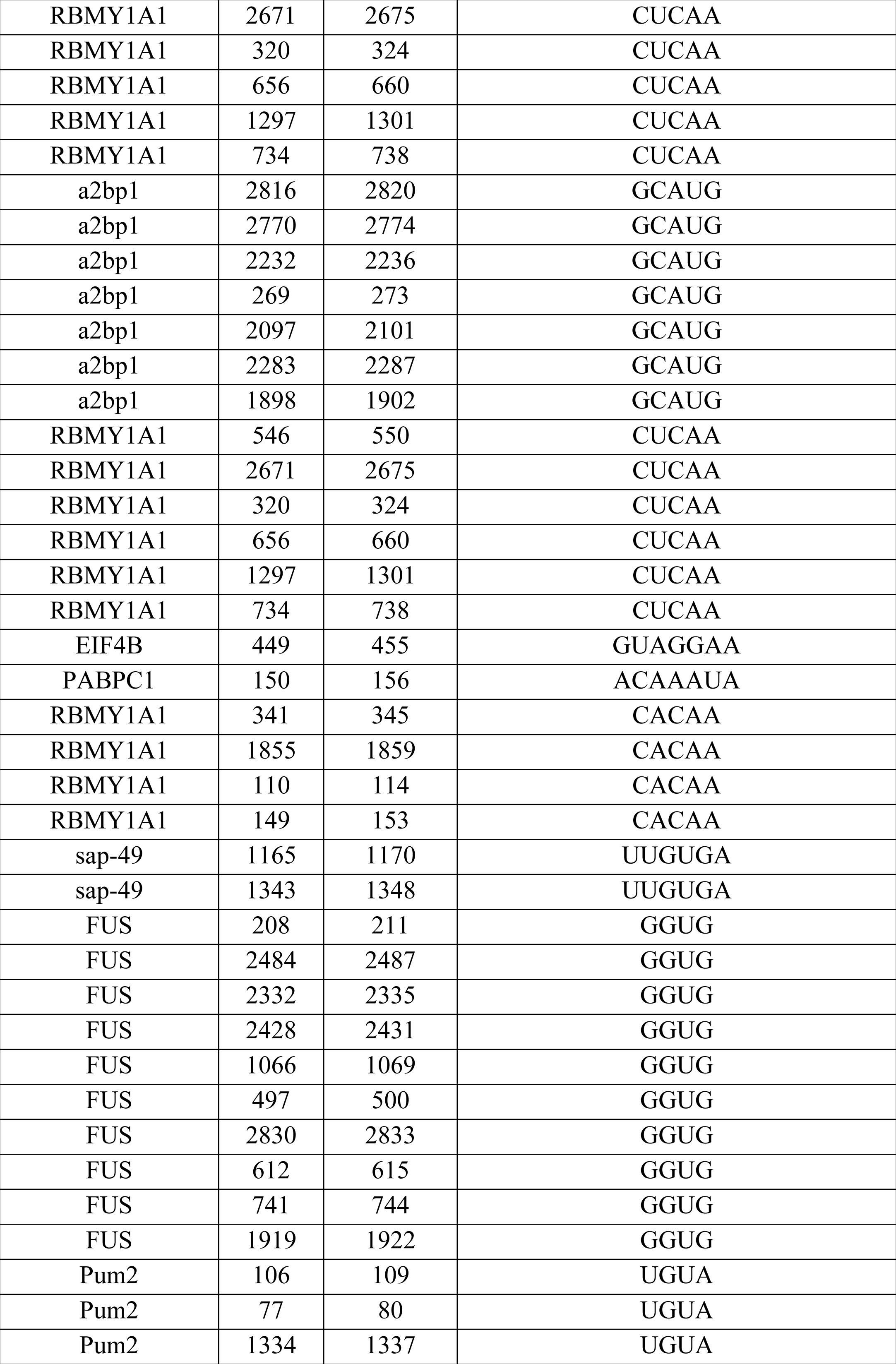
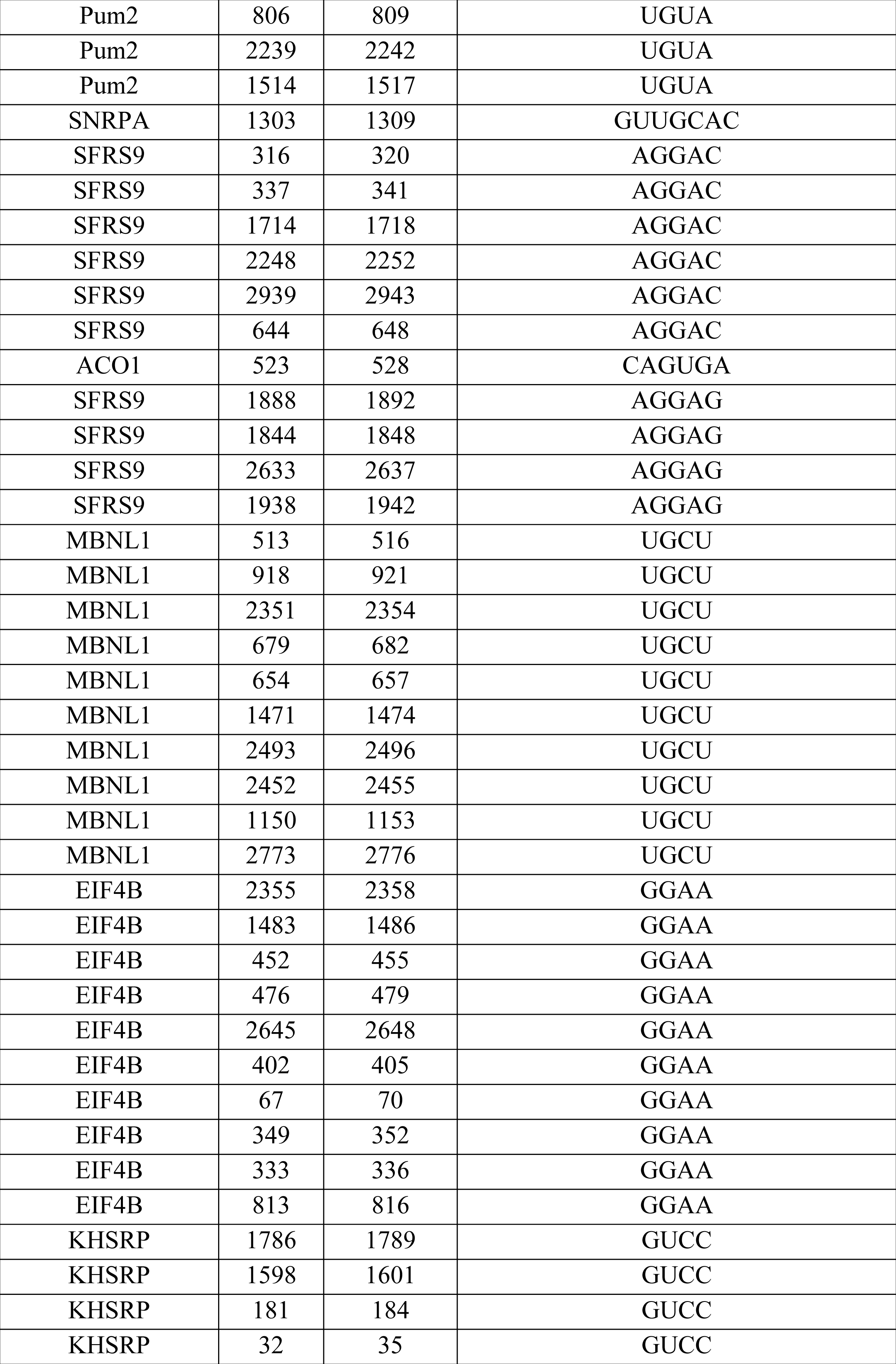
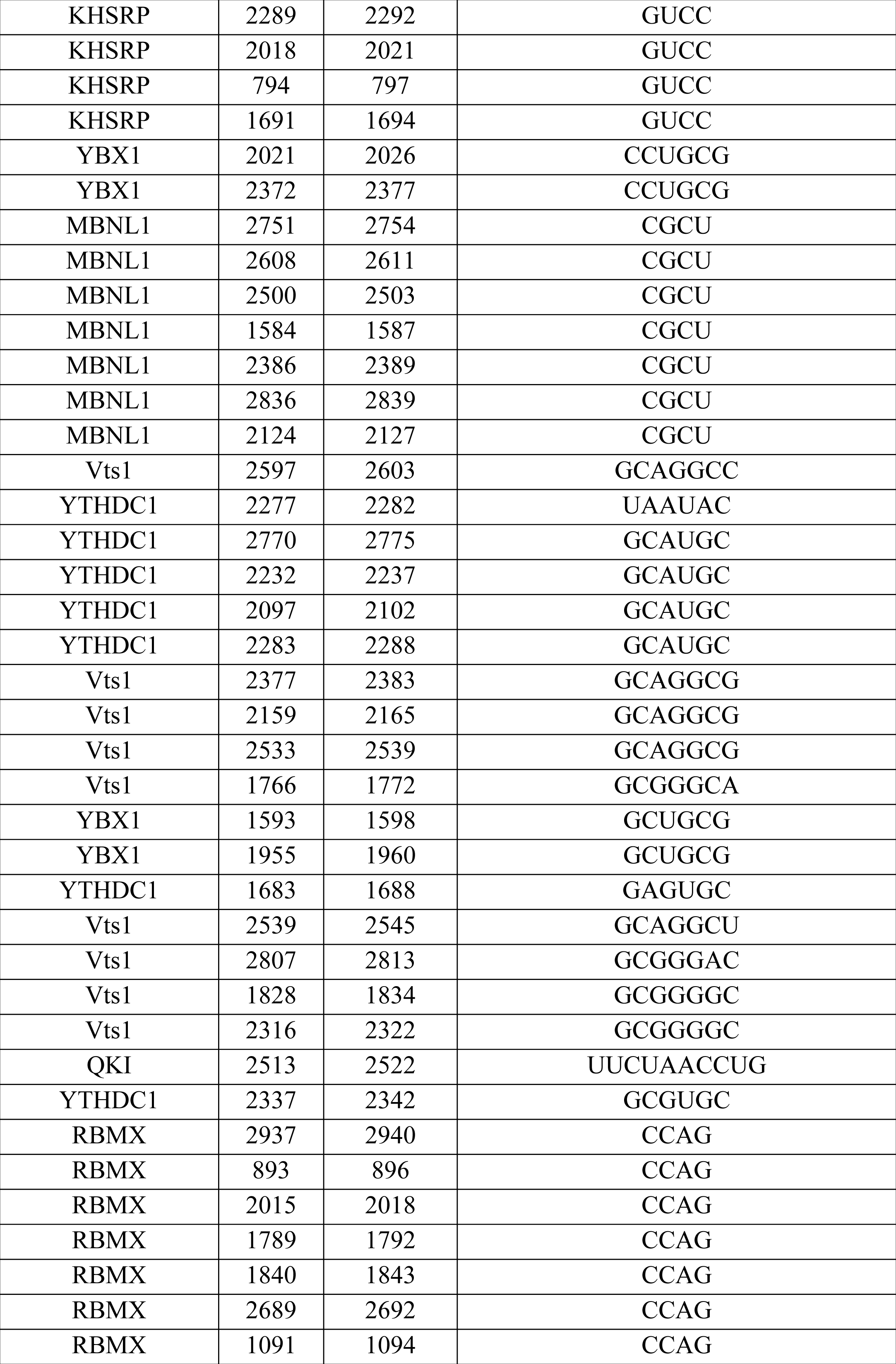
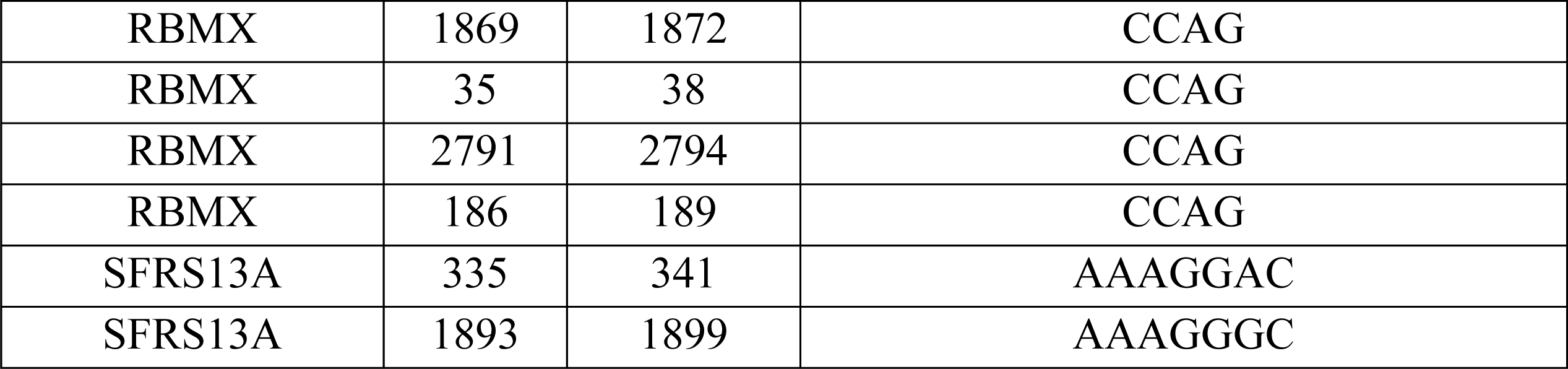
Predicted RNA binding proteins (RBPs) associated with the 5’ UTR of *Hnrnph1*. The 5’ UTR *Hnrnph1* sequence was loaded into RBPDB (61) where the sequence was scanned for putative sites for binding of RBPs. A list of RBPs that are predicted to bind to *Hnrnph1* and the binding sites on *Hnrnph1* associated with the RBP are shown. The start and end positions refer to the nucleotide position for the sequence in Supplemental Figure 1.

